# Power analysis for selection in allele-frequency time series

**DOI:** 10.64898/2026.11.23.768524

**Authors:** Yacine Ben Chehida, Inez Januszczak, Gavin Broad, Alyssa Paul, Ben Price, Ian Barnes

## Abstract

Allele frequencies tracked through time, from ancient DNA, museum collections or monitoring, offer a direct view of selection. Their value depends on whether the sampling design can distinguish change caused by selection from genetic drift and from the noise of sampling few individuals, often sequenced at low coverage. Researchers need to know how much power a design has before committing specimens and sequencing effort, which usually requires study-specific simulations. We present SPOT (Selection Power Over Time), a browser-based simulator that estimates the power to detect directional selection at a candidate locus from samples taken at several dates. Users describe the sampling design (dates, number of individuals, sequencing depth) and a biological scenario (inheritance system, generation time, effective population size, starting allele frequency, selection coefficient and dominance). Within seconds, SPOT simulates selection while accounting for sampling, genotype calling and drift, and sets the detection threshold from simulations without selection. Its sample-size planner shows how many individuals are needed to reach a target power. Conventional approaches, which model sampling error but not drift, overestimate power when drift is strong: in a museum insect case study, a threshold ignoring drift reported 69% power but called 10% of neutral loci selected instead of 5%; the drift-aware threshold gave 55%. SPOT agreed with the individual-based simulator SLiM in allele-frequency distributions, power and false-positive rates. Case studies of museum insects and human ancient DNA show how SPOT identifies detectable selection coefficients and helps allocate specimens and sequencing effort. SPOT is open source and runs without installation.

## 1. Introduction

Samples collected at different times provide a direct record of allele-frequency change. Ancient DNA, museum and herbarium collections, archived samples and long-term monitoring make such time series available for a growing range of species (Dehasque et al., 2020; Clark et al., 2025), and several methods test for, or estimate, selection from them (Mathieson & McVean, 2013; Feder et al., 2014; Foll et al., 2015; Ferrer-Admetlla et al., 2016; Schraiber et al., 2016; Barata et al., 2023; He et al., 2023; Cheng & Steinrücken, 2025). Before these methods are applied, or before material is sampled and sequenced, it is useful to know the power of the design: the probability that the available samples detect selection of plausible strength when it acts. Temporal samples are often small and unevenly spaced, and historical material is usually sequenced at low coverage, so the answer is not obvious.

In a time series, selection is detected as a change in allele frequency between sampling dates that is larger, or more consistent, than expected by chance (Feder et al., 2014). Two sources of chance contribute to the observed change. Sampling error arises because only a finite number of individuals is collected and genotyped at each date, so each sample gives only an estimate of the population’s allele frequency. Genetic drift arises because the population’s allele frequency itself changes at random from one generation to the next (Wright, 1931). Drift affects the detection of selection in two ways. It makes the trajectory of a favoured allele variable, so that an allele starting at low frequency can be lost before its advantage becomes visible. It also changes allele frequencies at neutral loci, and a putative signal of selection must be judged against these changes. Conventional tests that treat sampling as the only source of variation therefore reject neutral loci too often when drift is substantial (Waples, 1989; Wiberg et al., 2017; Spitzer et al., 2020), and a power calculation based on the same assumption overstates what a design can achieve. Adding individuals at each date reduces sampling error but not drift.

Accounting for drift is most practical by simulation. A realistic power analysis must include every source of chance (drift, sampling and genotyping error) and use a test statistic and a significance threshold calibrated under drift, evaluated over many simulations of each scenario. General forward simulators such as SLiM (Haller et al., 2026) and simuPOP (Peng & Kimmel, 2005) are flexible enough to do all of this, but only through study-specific code (Pinsky et al., 2021), which requires programming expertise and long computations, sometimes on high-performance computing clusters. Simulation studies of sampling designs exist for particular settings, including human ancient DNA time series (Anchieri et al., 2026), epidemic-driven selection (Santander & Moltke, 2025) and evolve-and-resequence experiments (Kofler & Schlötterer, 2014; Vlachos & Kofler, 2018). We are not aware of a general tool that estimates power for an arbitrary temporal sampling design while accounting for drift, sampling and sequencing depth.

Here we present SPOT (Selection Power Over Time), a browser-based simulator that estimates this power. We (i) describe its model and implementation; (ii) evaluate the false-positive rate of its threshold; (iii) compare its results with theory and with independent SLiM simulations; (iv) measure how much its power estimates vary between runs; (v) measure its speed; and (vi) test the accuracy of the rescaling procedure (Hoggart et al., 2007; Uricchio & Hernandez, 2014), a shortcut used for studies that span tens of thousands of generations, in which SPOT simulates several generations in one step to remain fast. Finally, (vii) we illustrate its use with two hypothetical case studies.

## 2. Materials and Methods

### 2.1. Principle of the method

A temporal study of selection samples a population at two or more dates, genotypes the sampled individuals at a candidate locus, and asks whether one allele (suspected to be under selection and which we call the favoured allele) became more frequent than expected by chance. The power of such a study is the probability that it detects selection when selection of a given strength acts on this allele. SPOT aims to estimate this power before any sample is collected or sequenced, or, when some samples already exist, to assess whether collecting or sequencing more would increase it.

Power depends on two kinds of input. The *design* is what the researcher chooses: the sampling dates, the number of individuals at each date, their sex and relatedness, and the sequencing depth. The *scenario* is the biology of the species, which is usually only partly known: inheritance system, generation time, effective population size (*N*_e_) and its changes, migration, starting frequency of the favoured allele, strength and dominance of selection, and the date at which selection starts. In SPOT, the user sets both (Table 1).

**Table 1.** Steps of one simulation.

| **Step** | **Description** | **User settings** |
|---|---|---|
| Initialisation | Frequency of the favoured allele *p*<sub>0</sub> in both sexes (in all individuals in haploids) at the first sampling date. | p<sub>0</sub> |
| Reproduction | Offspring genotype frequencies from those of the breeding adults: random union of gametes (haplodiploid males from their mother only; clonal reproduction in haploids). | Inheritance system |
| Selection | Viability selection with fitness 1, 1 + hs, 1 + s (diploid) or 1, 1 + s (haploid individuals), from the generation after the onset year; in males, *s* can be multiplied by a chosen factor. | s, h, selection in males, onset year |
| Sampling | Individuals drawn at each date from post-selection genotype frequencies. | Dates, individuals per date, probability of sampling a male, siblings per family |
| Genotyping | Poisson read depth, binomial allele reads with error, maximum-likelihood, majority or pseudo-haploid calls, minimum depth. | Mean depths, error rates, minimum depth; or perfect genotypes |
| Drift | Binomial draw of the gene copies of the next breeding adults (females and males; all individuals in haploids). | *N*<sub>f</sub>, *N*<sub>m</sub>; population change; *N*<sub>e</sub> per interval |
| Migration | Replacement of a fraction of breeding adults by migrants. | Migration rate, source frequency |
| Scoring | Trend statistic Z (all dates; first and last dates). | – |
| Power | Threshold: value of Z exceeded by a proportion *α*/*L* of neutral simulations; power: proportion of selection simulations above it. | *α*, *L*, numbers of simulations, seed |

Series longer than 2,000 generations may be rescaled (Section 2.6).

SPOT estimates power by simulating the planned study many times. A simulation follows the allele frequency of the population, generation by generation, from the first to the last sampling date. At each sampling date, it samples individuals and assigns them genotypes, as a sequencing study would. Each simulation therefore produces one simulated data set, the number of copies of each allele observed at each sampling date. Each data set differs because sampling, drift and genotype calls are random.

SPOT summarises each simulated data set by one number, the score *Z* (Cochran, 1954; Armitage, 1955), which measures how strongly the frequency of the favoured allele rises across the samples, from the first to the last sampling date (Section 2.5). *Z* is close to zero when the frequency does not change and large when it rises steadily. Because of these random processes, *Z* varies from one simulation to the next even without selection. SPOT uses this variation to set a threshold on *Z*, and then measures power, in two steps.

#### Step 1: setting the threshold with neutral simulations

SPOT runs neutral simulations (2,000 by default), which include the same drift, sampling and genotyping as the study, but no selection (selection coefficient *s*=0). Their values of *Z* show how large *Z* can become by chance, without selection. SPOT sets the threshold at the value of *Z* exceeded by 5% of the neutral simulations (more generally, by a proportion *α*, the significance level; Section 2.5 and Supporting Information S1.10).

#### Step 2: measuring power with selection simulations

SPOT also runs selection simulations (1,000 by default), which include the same drift, sampling and genotyping as the neutral simulations, plus selection: the favoured allele has a positive selection coefficient *s* chosen by the user. Power is the proportion of selection simulations whose *Z* exceeds the threshold set in step 1. Figure 1b illustrates the two steps for one design.

**Figure 1.**
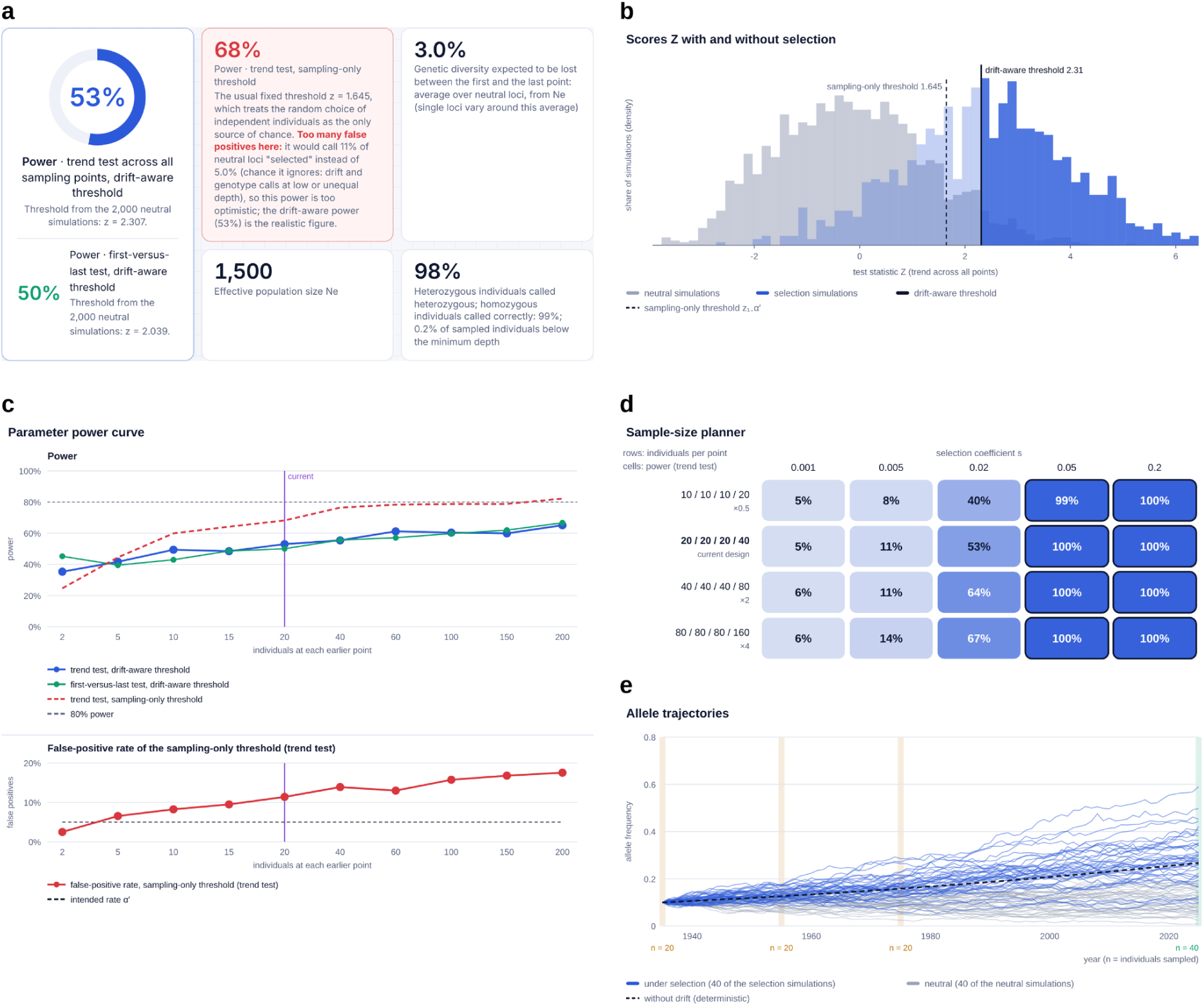
The SPOT interface for the museum insect case study (Table 3) with *p*_0_=0.1 and *s*=0.02. (a) Summary of results: power of the trend test with the drift-aware threshold (main tile; first-versus-last test below), power with the sampling-only threshold and its false-positive rate, expected loss of genetic diversity, *N*_e_ and genotype calls. (b) Distribution of the score *Z* in neutral simulations (grey) and selection simulations (blue); solid line, drift-aware threshold (2.31); dashed line, sampling-only threshold (1.645). (c) Power curve for the number of individuals at each historical date (top) and false-positive rate of the sampling-only threshold (bottom). (d) Sample-size planner: power for the design scaled by 0.5 to 4 (rows) across selection coefficients (columns); outlined cells reach 80%. (e) Allele-frequency trajectories of 40 selection simulations (blue) and 40 neutral simulations (grey) between the first and last sampling dates; dashed line, trajectory without drift; shaded bands, sampling dates, with the number of individuals sampled.

The threshold sets the risk of a false positive. A locus at which no selection acts behaves like a neutral simulation, so it exceeds the threshold, and is wrongly declared under selection, with a probability *α*: this is the false-positive rate of the test. A locus under selection of strength *s* behaves like a selection simulation, so it exceeds the threshold, and is correctly declared under selection, with a probability equal to the power. We call this threshold the drift-aware threshold, in this article and in the SPOT interface, because it is set from neutral simulations that include drift. For comparison, SPOT also applies to the same simulations the fixed threshold of tests that treat sampling as the only source of chance (Waples, 1989; Spitzer et al., 2020); we call it the sampling-only threshold (Section 2.5).

### 2.2. Population model

Each simulation follows one locus with two alleles in a single population with random mating (clonal reproduction in haploids) and non-overlapping generations (Wright–Fisher model; Wright, 1931). At the first sampling date, the favoured allele has the starting frequency *p*_0_ (“Allele frequency at the first point (p₀)” in the interface).

#### Time

SPOT counts time in generations. The user enters the sampling dates in calendar years and the generation time in years (“Years per generation”). The generation of each sampling date is the number of years since the first date divided by the generation time, rounded to the nearest whole number.

#### Inheritance systems

SPOT offers three inheritance systems: diploid, haplodiploid and haploid. There are no sexes in haploids.

#### How a generation is simulated

SPOT does not simulate individual organisms; it follows the frequency of the favoured allele in breeding females and males (in haploids, in all breeding individuals). Each generation goes through the five steps below, in this order.

1. Reproduction. The breeding adults mate at random (haploids reproduce clonally) and produce offspring. SPOT assumes that the pool of offspring is large enough that their genotype frequencies are exactly those expected from the allele frequencies of the breeding adults (Table 1), and calculates these frequencies exactly. Drift enters only at step 4, when a limited number of these offspring become the breeding adults of the next generation, as is standard in Wright–Fisher models.
2. Selection. Offspring survive in proportion to their fitness. Diploid offspring (including haplodiploid females) with zero, one and two copies of the favoured allele have fitness 1, 1 + *hs* and 1 + *s*, where *s* is the selection coefficient (“Selection coefficient s”) and *h* the dominance coefficient (“Dominance in diploid individuals”; *h*=0, recessive; 0.5, additive; 1, dominant). Haploid individuals, including haplodiploid males, have fitness 1 without the favoured allele and 1 + *s* with it. The setting “Selection in males (× s)” multiplies *s* in males, so that selection can be weaker, absent or stronger in males. Selection acts from the generation after the year set in “Selection starts in year”. By default, this is the first sampling date, so the first sample is taken before selection has acted.
3. Sampling. At a sampling date, individuals are sampled from the genotype frequencies of the offspring after selection (for siblings, see Section 2.3).
4. Drift. The breeding adults of the next generation are drawn at random from the offspring. For the *N*_f_ breeding females (“Breeding females per generation (*Nf*)”), the number of their 2*N*_f_ gene copies that carry the favoured allele is drawn from a binomial distribution whose probability is the frequency of this allele in female offspring after selection. The *N*_m_ breeding males (“Breeding males per generation (*Nm*)”) are drawn in the same way from male offspring, with 2*N*_m_ copies in diploids and *N*_m_ in haplodiploids. In haploids, the *N* breeding individuals (“Breeding individuals per generation (*N*)”) carry *N* copies. Drift can be switched off with “No drift (infinite population)”; the threshold is still set from neutral simulations, which then vary only through sampling and genotype calls (Section 2.5).
5. Migration. Migration (“Migrants per generation, *m*”) mixes the breeding adults with migrants from a source population whose allele frequency stays constant (“Allele frequency in the migrant source”; by default *p*_0_): the allele frequency of the breeding adults becomes a weighted mean of its previous value and that of the source, without chance.

#### Effective population size

The amount of drift is summarised by *N*_e_, which SPOT computes from the breeding numbers: *N*_e_=4*N*_f_*N*_m_/(*N*_f_ + *N*_m_) in diploids (Wright, 1931), *N*_e_=9*N*_f_*N*_m_/(4*N*_m_ + 2*N*_f_) in haplodiploids (Hedrick & Parker, 1997) and *N*_e_=*N* in haploids. Alternatively, *N*_e_ can be entered directly (“Enter Ne directly”). SPOT also reports the expected proportion of neutral genetic diversity, 2*p*(1 − *p*), lost between the first and last sampling dates, calculated as (Wright, 1931):

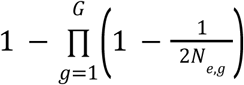

where *G* is the number of generations between these dates and *N*_e,*g*_ the value of *N*_e_ in generation *g*. In haploids, 1/(2*N*_e,*g*_) is replaced by 1/*N_g_*. With migration (rate *m*), each generation, a proportion 1 − (1−*m*)² of this diversity is restored to its initial value (Wright, 1931; S1.13).

*Changes in N*_e_ *over time.* Breeding numbers can (i) stay constant; (ii) change exponentially from the first to the last sampling date, by a chosen ratio of final to initial numbers (“Ne at the last point relative to the first”); or (iii) take a chosen *N*_e_ in each interval between consecutive sampling dates (“Set Ne separately for each interval”; details in S1.5).

### 2.3. Sampling of individuals

Three settings control the sampling step, in neutral and selection simulations alike: the number of individuals sampled at each sampling date, their sex and their relatedness. SPOT follows genotype frequencies rather than individuals: the genotype of each sampled individual is drawn at random from the genotype frequencies of the offspring of that generation, after selection when selection acts.

#### Sex of sampled individuals

In diploids and haplodiploids, sampled individuals are females by default; with “Males in the sample”, each sampled individual is instead a male with the chosen probability. A haplodiploid male carries one gene copy instead of two, and therefore contributes half as many gene copies to the sample as a female. In diploids, the sex of unrelated sampled individuals affects power only when selection differs between the sexes (“Selection in males (× s)” different from 1).

#### Related individuals

By default, sampled individuals are unrelated: each is drawn independently of the others. They can instead be sampled in groups of full siblings, which share both parents (haplodiploid brothers share only their mother), by setting “Sampled individuals per family” above 1. This represents, for example, close relatives in human ancient DNA or nestmates of a singly mated queen in social insects. Because siblings share their parents, their genotypes are similar, and a group of siblings carries less information on the allele frequency of the population than the same number of unrelated individuals. The more siblings per family, the fewer independent families a sample contains. For each family, SPOT draws the mother’s and father’s genotypes from the genotype frequencies of female and male offspring after selection, and those of the siblings from their parents following Mendel’s rules, without further selection (S1.7). In haploids, this setting has no effect.

### 2.4. Sequencing and genotype calls

Once individuals are sampled, SPOT assigns them genotypes in one of two ways. With “Perfect genotypes (ignore sequencing)”, the true genotype of each sampled individual is used. Otherwise, SPOT simulates the sequencing reads of each sampled individual at the locus and calls its genotype from these reads. Simulated sequencing reproduces the two main consequences of low sequencing depth: missing genotypes and wrong genotype calls. The per-read error rate represents all causes of reads showing the wrong allele, such as sequencing errors and post-mortem damage in ancient DNA, as a single probability that is the same for both alleles, without modelling their mechanisms.

#### Read depth

For each sampled individual, the number of reads covering the locus, the read depth, is drawn at random from a Poisson distribution whose mean, the mean depth, is set by the user. Two mean depths are set: one for all sampling dates except the last (“Mean depth, earlier points”), and one for the last sampling date (“Mean depth, last point”). This allows, for example, historical samples sequenced at low depth to be combined with recent samples sequenced at a higher depth.

#### Read errors

Each read comes from one of the gene copies of the individual (chosen at random in diploid individuals) and shows the wrong allele with probability ε, which can differ between the earlier dates and the last date (“Per-read error rate, earlier points” and “Per-read error rate, last point”), because historical samples often carry more errors, due to post-mortem DNA damage (Briggs et al., 2007).

#### Genotype calls

Individuals with fewer reads than “Minimum depth to call a genotype” receive no genotype and are left out. Because read depth varies among individuals, some are left out even when the mean depth is above this minimum, so not all sampled individuals contribute to *Z*. For the remaining individuals, SPOT calls the genotype that best explains the reads, that is, the genotype under which the observed numbers of reads of each allele are most probable (maximum-likelihood calling; Nielsen et al., 2011; S1.8). For haploid individuals, the allele shown by most reads is called (ties are resolved at random).

#### Pseudo-haploid calls

SPOT can also use pseudo-haploid calls, an approach common in ancient DNA (Mallick et al., 2024) in which one random read per individual counts as one gene copy (“Genotype calls”: “Pseudo-haploid”; S1.8, S7.3).

### 2.5. Score and significance thresholds

#### Score

The score *Z* is computed from the called gene copies of each simulated data set. *Z* is the Cochran–Armitage trend statistic (Cochran, 1954; Armitage, 1955), a standard test assessing whether the frequency of the favoured allele in the samples changes linearly with time, counted in generations (details in S1.9). *Z* is positive when the favoured allele is more frequent in later samples than in earlier ones, and increases with the size of this rise and with the number of gene copies sampled. The test is one-sided, as only an increase of the favoured allele counts as evidence of selection.

#### Two versions of the score

SPOT computes *Z* in two versions: with all sampling dates (the trend test) and with the first and last sampling dates only (the first-versus-last test). Each version has its own threshold, set as in Section 2.1. With only two sampling dates, the two versions are identical.

#### Significance level and number of simulations

The significance level *α* is set by the user (“Significance level α”; 0.05 by default). When *L* candidate loci are tested (“Candidate loci tested, L (each tested at α′ = α/L)”), each locus is tested at the level *α*′=*α*/*L* (Bonferroni correction; Dunn, 1961), so that the probability of any false positive among *L* neutral loci stays at or below *α*; the threshold and power are then computed for one locus with the chosen scenario, tested at level *α*′. The threshold is set so that a proportion *α*′ of the neutral simulations exceed it (S1.10). By default, SPOT runs 1,000 selection simulations (“Selection simulations (at least twice as many neutral)”) and twice as many neutral simulations. When *α*′ is small, more neutral simulations are needed to place the threshold reliably, and SPOT increases their number to 10/*α*′, so that about ten neutral values are expected above the threshold, up to 200,000 (that is, up to 1,000 loci at *α*=0.05); beyond this, it fits a normal distribution to the neutral values of *Z* and marks the threshold as unreliable (S1.12).

#### Sampling-only threshold

SPOT also reports the results of a standard test whose threshold comes from a formula rather than from simulations. This test assumes that the sampled gene copies are independent and that their sampling is the only source of chance. Under these assumptions, *Z* approximately follows a standard normal distribution without selection, so the test uses a fixed threshold, *z*_1−*α*′_, the 1 − *α*′ quantile of the standard normal distribution (1.645 for *α*′=0.05 and one locus). SPOT applies this threshold to the same neutral and selection simulations and reports, for the trend test, its actual false-positive rate and the power it would report.

### 2.6. Rescaling of long time series

Studies of organisms with short generations, such as bacteria or viruses, can span tens of thousands of generations (Good et al., 2017). Simulating every generation would make such studies slow, so SPOT simulates a smaller population in which each simulated generation stands for *λ* real generations (the rescaling factor), a shortcut common in forward-in-time simulations (Hoggart et al., 2007; Uricchio & Hernandez, 2014) that approximately reproduces the total amount of drift and effect of selection over the study period.

When the first and last sampling dates are more than 2,000 generations apart, SPOT divides two quantities by *λ*: *N*_e_ and the generation numbers of the sampling dates. Because the variance of drift per generation is inversely proportional to *N*_e_, dividing both the number of generations and *N*_e_ by *λ* leaves the total drift unchanged. The selection coefficient becomes (1 + *s*)*^λ^* − 1, the fitness advantage accumulated over *λ* generations by a haploid or homozygous carrier (*h* is unchanged). For small *s*, this is close to *λs*, as in the usual rescaling (Hoggart et al., 2007), so *N*_e_*s*, which sets the strength of selection relative to drift, is kept. The migrant fraction *m* becomes 1 − (1 − *m*)*^λ^*. Details, the limits of rescaling and its validation are given in Section S6.

### 2.7. Implementation

The model described above is implemented in SPOT, an HTML file written in JavaScript that runs locally in a web browser, without installation. The interface (Figure 1) offers six example scenarios. Its outputs are listed in Table 2. Besides the power of the design, they include the simulated allele-frequency trajectories (Figure 1e); a power curve, which shows how power changes when one setting is varied while all others are kept fixed (Figure 1c); a sample-size planner (Figure 1d), which shows power for designs with 0.5, 1, 2 and 4 times as many individuals over a range of one setting, such as the selection coefficient. To remain fast, outputs that compute several designs, such as the power curve and the sample-size planner (20 designs), run at most 1,000 selection simulations per design. Their neutral simulations follow the rule of Section 2.5, but are capped at 20,000 per design instead of 200,000. Their thresholds therefore come from the normal fit with more than 100 loci at *α*=0.05, instead of more than 1,000 loci for a single calculation (Section 2.5). The simulation code is also distributed as a separate file (sim.js) that runs in Node.js, outside the browser.

**Table 2.** Outputs of SPOT.

| **Output** | **Description** |
|---|---|
| Power | Trend test and first-versus-last test, with the drift-aware threshold. |
| Sampling-only threshold | Proportion of neutral simulations above the fixed threshold z<sub>1-</sub>α′ (actual false-positive rate) and power this threshold would report. |
| Scenario summary | *N*<sub>e</sub>, expected loss of neutral genetic diversity (with and without migration), proportions of heterozygous and homozygous individuals (or haploid gene copies) called correctly, proportion of individuals below the minimum depth; when above 50%, proportion of neutral simulations in which drift lost or fixed the allele by the last date. |
| Charts | Simulated trajectories with and without selection, and the expected frequencies without drift; distributions of Z with both thresholds. |
| Power curve | Power over a range of one parameter (e.g. s, h, p<sub>0</sub>, *N*<sub>e</sub>, onset year, individuals per date, depth). |
| Sample-size planner | Power for the design scaled by 0.5, 1, 2 and 4, over a range of one parameter. |
| Sampling of males (haplodiploids) | Power when sampling females, males or both. |
| Reproducibility | Seed; settings in the page address or a file; CSV and image export; sim.js for scripted use. |

### 2.8. Evaluation

We evaluated SPOT in four ways: the false-positive rate of the drift-aware threshold (Section 2.8.1); the agreement of its simulations with theoretical expectations (loss of genetic diversity under drift, including changes in *N*_e_ and migration, trajectories under selection with and without drift, exact power without drift, and the sampling of relatives; S3) and with the individual-based simulator SLiM (Section 2.8.2); the precision and speed of a single calculation (S5); and the effect of rescaling on power and on false-positive rate (S6).

All analyses used sim.js, with the trend test, *α*=0.05 and one locus unless stated otherwise. All analyses were performed on CropDiversity HPC (Percival-Alwyn et al., 2025). Each analysis used several test cases, each a complete set of settings (design and scenario); their full settings are given in Sections S2–S6. Except in the analyses of precision (S5) and rescaling (S6), all test cases were derived from one reference case: a haplodiploid insect with *N*_e_=1,500 and p₀=0.1, sampled in 1935, 1955, 1975 and 2025 (20, 20, 20 and 40 females; one generation per year, so 90 generations), which is also the museum insect case study of Section 2.9 (Table 3; Figure 1).

**Table 3.** Case study settings.

| **Setting** | Museum insect | **Human ancient DNA** | **Bacterial pathogen (S6)** |
|---|---|---|---|
| Inheritance | Haplodiploid | Diploid | Haploid |
| Sampling years | 1935, 1955, 1975, 2025 | 7000, 5000, 3000, 1000 BCE; 1000, 2000 CE | 1850, 1900, 1950, 2000, 2020 |
| Generation time | 1 year | 28 years | 0.01 year (100 generations per year) |
| Generations | 90 | 321 | 17,000 (simulated as 1,889 with rescaling, λ=9) |
| Individuals per date | 20, 20, 20, 40 females | 50 per ancient date, 100 present-day | 10, 10, 15, 20, 40 |
| *N*<sub>e</sub> | 1,500 (150, 300, 450, 750, 1,500, 3,000, 7,500, 30,000 and 150,000 in Figure 3b,c) | 10,000 | 100,000 |
| Selection coefficient *s* | 0.005, 0.01, 0.02, 0.03, 0.05, 0.075, 0.1, 0.15 and 0.2 (Figure 3a); 0.05, 0.02 and 0.015 for *p*<sub>0</sub>=0.01, 0.1 and 0.3 (Figure 3b,c, as in Figure 3d); 0.02 (Figure 1, Figure 3e); Figure 3d: see note | 0.001–0.1 (Figure S6a); 0.005 and 0.0075 (Figure S6b) | 0.00002–0.0005 per generation (Figure S5) |
| Starting frequency | 0.01, 0.1, 0.3 | 0.1 | 0.05 |
| Mean depth, earlier/ last date | 10×/15× (1–30× at all dates in Figure 3e) | 2×/20× (1–10× in Figure S6) | 10×/30× |
| Selection simulations | 1,000 (5,000 in Figure 3d,e) | 1,000 (5,000 in Figure S6b) | 5,000 |

All scenarios are hypothetical. h=0.5, per-read error rate 0.005 (human ancient DNA: 0.01), minimum depth 3 (human ancient DNA and Figure 3e: 1), individuals from different families, no migration, selection from the first date; twice as many neutral as selection simulations. Selection coefficients for Figure 3d: (p<sub>0</sub>, s)=(0.01, 0.05), (0.1, 0.02), (0.3, 0.015) at *N*<sub>e</sub>=1,500, and (0.01, 0.035), (0.1, 0.015), (0.3, 0.012) at 100,000.

#### 2.8.1. False-positive rate of the threshold

Power is meaningful only if the threshold has the intended false-positive rate. On the neutral simulations used to set a threshold, a proportion *α*′ lie above it by construction; whether the threshold keeps this false-positive rate must therefore be measured on other neutral simulations, generated independently. For each test case, we ran SPOT 20 times with different random seeds, obtaining 20 thresholds. We then ran 100,000 additional, independent neutral simulations with the same settings and applied each of the 20 thresholds to them, one by one: the proportion of these simulations above a threshold is its realised false-positive rate, which should be close to or below α′ if the threshold is correct. We applied the sampling-only threshold to the same simulations (S2).

We used 36 test cases (Tables S1 and S2). The first is the reference case. In 24 others, we changed its inheritance system and *N*_e_, to vary drift. In the remaining 11, we changed one other setting at a time: two or ten sampling dates, a starting frequency of 0.01 or 0.5, a mean depth of 2×, perfect genotypes, four siblings per family, a decline, a bottleneck, migration, or 200 individuals per date.

To check the threshold at the much smaller significance levels used when many loci are tested, we repeated this analysis for four of these 36 test cases: (1) the reference case, (2) the reference case with *p*_0_=0.01, (3) the haplodiploid test case with *N*_e_=300 and (4) the haplodiploid test case with *N*_e_=100,000, changing only the number of candidate loci (10, 100 and 1,000 instead of one). Their realised rates were measured on 100,000 independent neutral simulations with 10 loci, and on 1,000,000 with 100 and 1,000 loci (Table S3). In these four test cases, we also measured the realised rate of a threshold from a normal distribution fitted to 2,000 neutral simulations, to assess whether such a fit could replace the additional neutral simulations.

#### 2.8.2. Comparison with SLiM

SPOT is fast because it simplifies the simulation by following allele frequencies rather than individuals (S4). To test whether these simplifications give the same results as a simulation of individual organisms, we compared SPOT with SLiM 5.2 (Haller et al., 2026), a widely used forward simulator that models every individual. Both programs simulated the same 11 scenarios: six with selection, which differed in inheritance system, *N*_e_, starting frequency, selection coefficient and dominance, and five without selection (Table S5). All used the sampling design of the reference case with perfect genotypes, so this comparison concerns drift, selection and sampling, not genotyping. Each SLiM simulation recorded, at each sampling date, the number of copies of the favoured allele among the sampled females, the same quantity that SPOT simulates. We compared the simulations generated by SPOT and SLiM in four ways (S4):

1. *Allele frequencies.* The sampling dates of the reference case correspond to generations 0, 20, 40 and 90. Both programs start from *p*_0_ at generation 0, so for each scenario we compared the 2,000 SPOT and 2,000 SLiM simulations at generations 20, 40 and 90: the mean allele frequency among breeding females after selection (difference in standard errors) and, at generation 90, its whole distribution (two-sample Kolmogorov–Smirnov test).
2. *Power.* For each of the six scenarios with selection, we estimated power 20 times independently with each program. For both programs, *Z* was computed as in Section 2.5, using only the simulations of that program: its 2,000 neutral simulations set the threshold, and power was the proportion of its 2,000 selection simulations above this threshold. We then compared the mean and the standard deviation of the 20 estimates of SPOT with those of SLiM.
3. *False-positive rates.* To test whether a threshold set from SPOT’s simulations gives the same false-positive rate on SLiM’s simulations as on SPOT’s own, we applied a threshold set from 20,000 SPOT neutral simulations (ten times the default, for a more stable threshold), and the sampling-only threshold, to 10,000 SLiM neutral simulations of each neutral scenario and to 10,000 independent SPOT neutral simulations. If SPOT simulates drift as SLiM does, the SPOT threshold is expected to give about 5% false positives on SLiM’s simulations.
4. *Power with SPOT’s threshold.* To test whether the power reported by SPOT holds in populations simulated individual by individual, we applied a threshold set from 20,000 SPOT neutral simulations to the selection simulations of both programs in each of the 20 runs, and compared the mean power on SLiM’s and on SPOT’s simulations.

### 2.9. Case studies

We analysed two hypothetical case studies (Table 3; tables and figures in S7); a third, a bacterial pathogen simulated with rescaling, is presented in Section S6. Results are for the trend test with the drift-aware threshold, *α*=0.05 and one locus.

*Museum insect case study.* This case study is the reference case described in Section 2.8: a haplodiploid insect sampled from museum collections in 1935, 1955 and 1975 and in the field in 2025 (females only).

a. Which selection coefficients can the design detect? We computed power for *s* from 0.005 to 0.2 at three starting frequencies (*p*_0_=0.01, 0.1 and 0.3).
b. How does power depend on *N*_e_, and can more individuals compensate for strong drift? We studied three combinations of starting frequency and selection coefficient: *p*_0_*=*0.01 with *s*=0.05, *p*_0_*=*0.1 with *s*=0.02, and *p*_0_=0.3 with *s*=0.015. The selection coefficient differs between the three combinations so that each gives intermediate power (55–65%) with the reference samples at *N*_e_=1,500. For each combination, we varied *N*_e_ from 150 to 150,000 (Table 3), and computed power with the reference number of samples and with two and five times as many individuals at every sampling date.
c. Where is extra effort most useful? We compared the reference sampling (100 individuals) with: 60 more historical specimens (40 at each sampling date); 60 more modern specimens (100 in 2025); and the same 100 specimens concentrated at the first and last sampling dates (50 at each). We made this comparison at *N*_e_=1,500 and at *N*_e_=100,000, each with *p*_0_=0.01, 0.1 and 0.3, with 5,000 selection simulations per design, and with *s* chosen so that the reference sampling had intermediate power (settings in Table 3 and Table S9).
d. How much does sequencing depth matter? We sequenced the whole design at the same mean depth at all sampling dates, 1×, 2×, 5×, 10× or 30×, and compared it with perfect genotypes, at *N*_e_=1,500 and 100,000, with *p*_0_=0.1 and *s*=0.02. The minimum depth to call a genotype was 1. To represent post-mortem damage, the 1× and 2× designs were also run with per-read error rates of 0.02 and 0.05 instead of 0.005; the same error rate was used at all sampling dates. We also ran designs with 2× at the earlier dates and 30× at the last date, and the reverse, and the 1× and 2× designs with the default minimum depth of 3. Each design used 5,000 selection simulations.

#### Human ancient DNA

We used the “Human ancient DNA” example scenario of the application (Table 3): six sampling dates over 321 generations, 50 individuals per ancient date at 2× and 100 present-day individuals at 20×, kept in all designs.

a. How much does the depth of the ancient samples matter? We computed power with the ancient samples sequenced at 1×, 2×, 5× or 10×, or with perfect genotypes, for *s*=0.001, 0.002, 0.003, 0.005, 0.0075, 0.01, 0.02, 0.05 and 0.1, with 1,000 selection simulations per design.
b. With a fixed sequencing effort, is it better to sequence many individuals at low depth or few at higher depth? We compared designs with the same number of reads at the locus per ancient date (number of individuals×mean depth=100): 100 individuals at 1×, 50 at 2×, 25 at 4×, 20 at 5× or 10 at 10×. We used *s*=0.005 and 0.0075, which gave intermediate power, and 5,000 selection simulations per design.
c. Does power change with pseudo-haploid calls, an approach common in human ancient DNA, in which one read drawn at random from each individual counts as one gene copy? With the ancient samples at 1×, 2×, 5× or 10× and s=0.005 and 0.0075, we analysed the same simulated reads with SPOT’s genotype calls and with pseudo-haploid calls, each with a threshold from neutral simulations analysed the same way (5,000 selection simulations per design).

## 3. Results

### 3.1. False-positive rate of the threshold

The drift-aware threshold gave close to the intended false-positive rate in all 36 test cases: at a significance level of 5%, the mean realised rate of the 20 thresholds of a test case was 4.83– 5.27% for the trend test and 4.84–5.25% for the first-versus-last test (Figure 2a,b; Table 4; Tables S1, S2). Individual thresholds, each from one run of SPOT, gave 3.6–6.7% for the trend test and 3.5–7.3% for the first-versus-last test.

**Figure 2.**
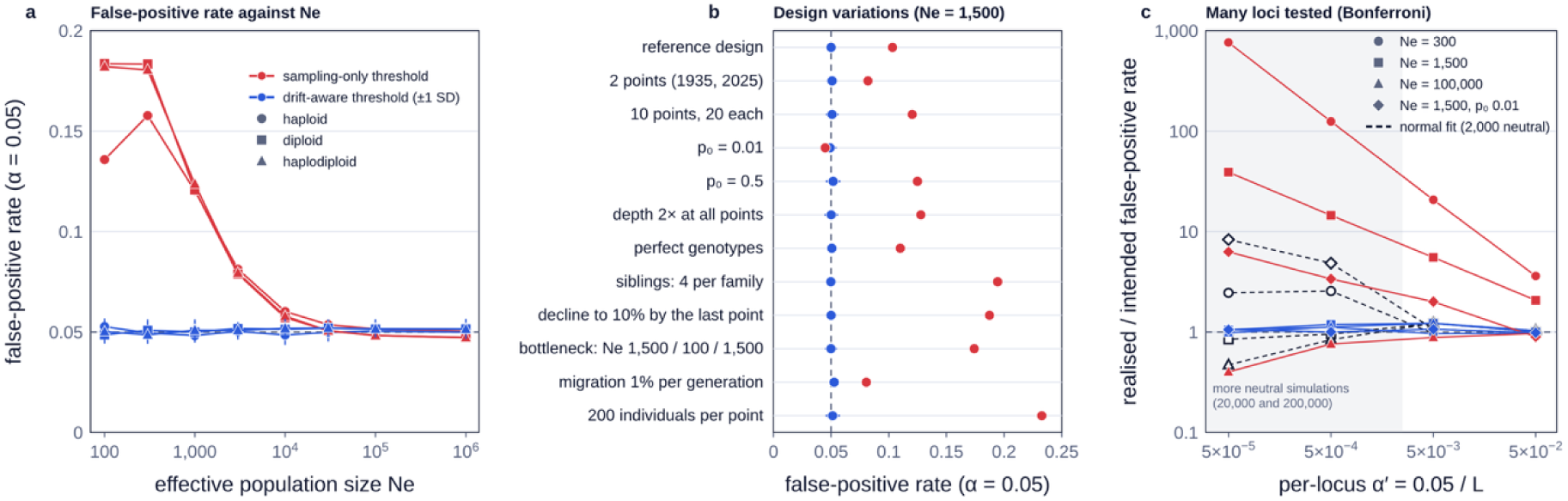
Realised false-positive rates of the trend test for an intended rate of 5%. Reference case: haplodiploid, sampled in 1935, 1955, 1975 and 2025 (20, 20, 20 and 40 females), mean depth 10× for the historical and 15× for the modern samples, *p*_0_=0.1. Blue, drift-aware threshold: mean realised rate (± SD in a and b) of 20 thresholds, each set by SPOT from its own neutral simulations (2,000 by default; up to 200,000 in c) and applied to 100,000 independent neutral simulations (1,000,000 for 100 and 1,000 loci in c); red, sampling-only threshold. (a) *N*_e_ and inheritance system (marker shape). (b) Reference case (*N*_e_=1,500) and versions of it in which one setting was changed at a time. (c) Ratio of realised to intended rate for *L* loci tested at *α*/*L*, for four test cases (marker shape); shaded, levels at which SPOT runs more neutral simulations (20,000 and 200,000); dashed black, threshold from a normal distribution fitted to 2,000 neutral simulations (Table S3).

**Table 4.** Summary of the evaluation.

| **Analysis** | Processes and conditions | **Main result** |
|---|---|---|
| False-positive rate | 36 test cases: reference design (90 generations) with three inheritance systems and *N*<sub>e</sub> 100–10<sup>6</sup>, or with one setting changed; drift, sampling and genotyping; 20 thresholds per test case (2,000 neutral simulations each), each applied to 100,000 independent neutral simulations | Drift-aware threshold: mean per test case 4.83–5.27% (trend test; single runs 3.6–6.7%) for an intended 5%; sampling-only threshold 4.5–23.3% (trend test) |
| Multiple loci | 4 haplodiploid test cases (*N*<sub>e</sub> 300, 1,500, 100,000; p<sub>0</sub> 0.1 or 0.01); *L*=10, 100, 1,000 (α′ down to 5 × 10<sup>−5</sup>); 100,000 or 1,000,000 independent neutral simulations | Realised/intended 0.96–1.22 (drift-aware threshold, trend test); sampling-only threshold up to 763 |
| SLiM 5.2 | 6 scenarios with and 5 without selection (three inheritance systems; *N*<sub>e</sub> 200–1,500); reference sampling design, perfect genotypes; drift, selection and sampling; 20 runs of 2,000 selection and 2,000 neutral simulations per program; 10,000 SLiM neutral simulations per neutral scenario for false positives; SPOT's threshold applied to both programs' selection simulations | Same allele-frequency distributions (KS p > 0.15); mean power differences −0.9 to +1.0 points (all within 2 SE); SPOT threshold on SLiM 4.9–5.5% false positives (4.9–5.7% on SPOT); sampling-only threshold 5.4–19.7% on both; power with SPOT's threshold on SLiM within 0.4 points of that on SPOT |
| Precision | 5 scenarios; 50 runs each with 500–5,000 selection simulations | SD of power 1.5–2.2 points (trend test) with 1,000 selection simulations, 0.6–1.1 with 5,000; one calculation 0.05–0.57 s (one locus), about 7 s with 1,000 loci |
| Rescaling | 120 haploid, diploid and haplodiploid test cases; 5,000–60,000 generations; λ=3–30; power of the trend test, full against rescaled simulations; false-positive rate of rescaled thresholds on 50,000 full neutral simulations (9 test cases) | SD of differences 2.4 points (107 test cases with power below 100%); false positives 4.7–5.2%; median speed-up 4.4 (max 28) |

Details in Sections S2–S6.

The sampling-only threshold, in contrast, usually gave more false positives than intended. For the trend test, the difference was particularly large when the samples contained chance variation not due to sampling, that is, drift, relatedness or low-depth genotype calls (Waples, 1989): its rate was 13.6–18.4% at *N*_e_=100–300, 18.7% with a decline of the population to 10%, 19.4% with four siblings sampled per family and 12.8% with a mean depth of 2× at all dates, compared with 10.3% for the reference case. With 200 individuals per date, it reached 23.3%, because larger samples reduce sampling error but not drift. Where drift was negligible, at *N*_e_ of 10^5^ or more, the rate was close to 5%. For a rare allele (*p*_0_=0.01), it gave fewer false positives than intended: 4.5% for the trend test and 1.7% for the first-versus-last test (Figure 2b).

When many loci were tested, the drift-aware threshold kept its intended rate: for the trend test, the mean ratio of realised to intended rate was 0.96–1.22 at all levels down to *α*′=5×10^−5^ (*i.e.* 1,000 loci; Figure 2c; Table S3). A normal distribution fitted to 2,000 neutral simulations could not replace the additional neutral simulations: with 100 and 1,000 loci, the resulting threshold gave 0.5–8.3 times the intended rate for the trend test, and far fewer false positives than intended for the first-versus-last test (Figure 2c). The sampling-only threshold gave up to 763 times the intended rate with strong drift (*N*_e_=300, 1,000 loci), but fewer false positives than intended with weak drift (*N*_e_=100,000).

### 3.2. Comparison with SLiM

The distributions of the population allele frequency were similar between SPOT and SLiM: mean frequencies at all generations differed by at most 2.2 standard errors, and the whole distributions at generation 90 showed no significant difference in any of the 11 scenarios (Kolmogorov–Smirnov *D* ≤ 0.035, *p* > 0.15). In each of the six scenarios with selection, power was estimated 20 times with each program. The mean of the 20 estimates differed between the programs by −0.9 to +1.0 percentage points (hereafter, points), never by more than two standard errors. The 20 estimates also varied by similar amounts in the two programs: their standard deviation was 1.3–2.4 points (Figure S2a; Table S5).

The two thresholds behaved on SLiM’s simulations as on SPOT’s. SPOT’s drift-aware threshold gave 4.9–5.5% false positives on SLiM’s neutral simulations, compared with 4.9– 5.7% on independent SPOT neutral simulations (Table S6). The sampling-only threshold gave 5.4–19.7% on the simulations of both programs, which shows that its higher false-positive rate is not a property of SPOT’s model. SPOT’s threshold also gave the same power on SLiM’s populations with selection as on SPOT’s: mean differences ranged from −0.4 to +0.3 points over 20 runs (Table S5). SPOT also agreed with the theoretical expectations for drift, for selection and, without drift, for power (S3; Table S4; Figure S1).

### 3.3. Precision, speed and rescaling

With the default 1,000 selection simulations, power varied between runs with a standard deviation of 1.5–2.2 points for the trend test (0.6–1.1 with 5,000 selection simulations), and a complete calculation took less than 1 s with one locus and about 7 s with 1,000 candidate loci (S5). For a final estimate, we therefore recommend 5,000 selection simulations.

Rescaling did not bias power on average. We compared power with rescaling and with full simulations in 120 test cases spanning 5,000 to 60,000 generations. In 13, power was 100% with both. In the other 107, power with rescaling was higher in 50, lower in 53 and equal in four, and the differences had a standard deviation of 2.4 points (1.9 without one haplodiploid test case that differed by 14.7 points, but repeated runs of this test case varied by more than ten points with either method; S6). Thresholds set from rescaled neutral simulations gave a mean of 4.7–5.2% false positives on neutral simulations without rescaling (nine test cases). Rescaling reduced run time by a median factor of 4.4, and up to 28 (Figure S4; Table S8).

### 3.4. Case studies

*Museum insect case study.* The reference case illustrates the difference between the two thresholds (Figure 1; *p*_0_=0.1, *s*=0.02). Over 20 runs of 5,000 selection simulations, the power of the trend test was 55% on average (53% in the run of Figure 1, with a drift-aware threshold of 2.31). The sampling-only threshold (1.645) was exceeded by 10% of the neutral simulations and would have reported a power of 69%.

The selection coefficient that this design can detect depended strongly on the starting frequency of the favoured allele: 80% power required *s* of about 0.065 for *p*_0_=0.01, 0.027 for 0.1 and 0.022 for 0.3, by linear interpolation in *s* between simulated values (Figure 3a).

**Figure 3.**
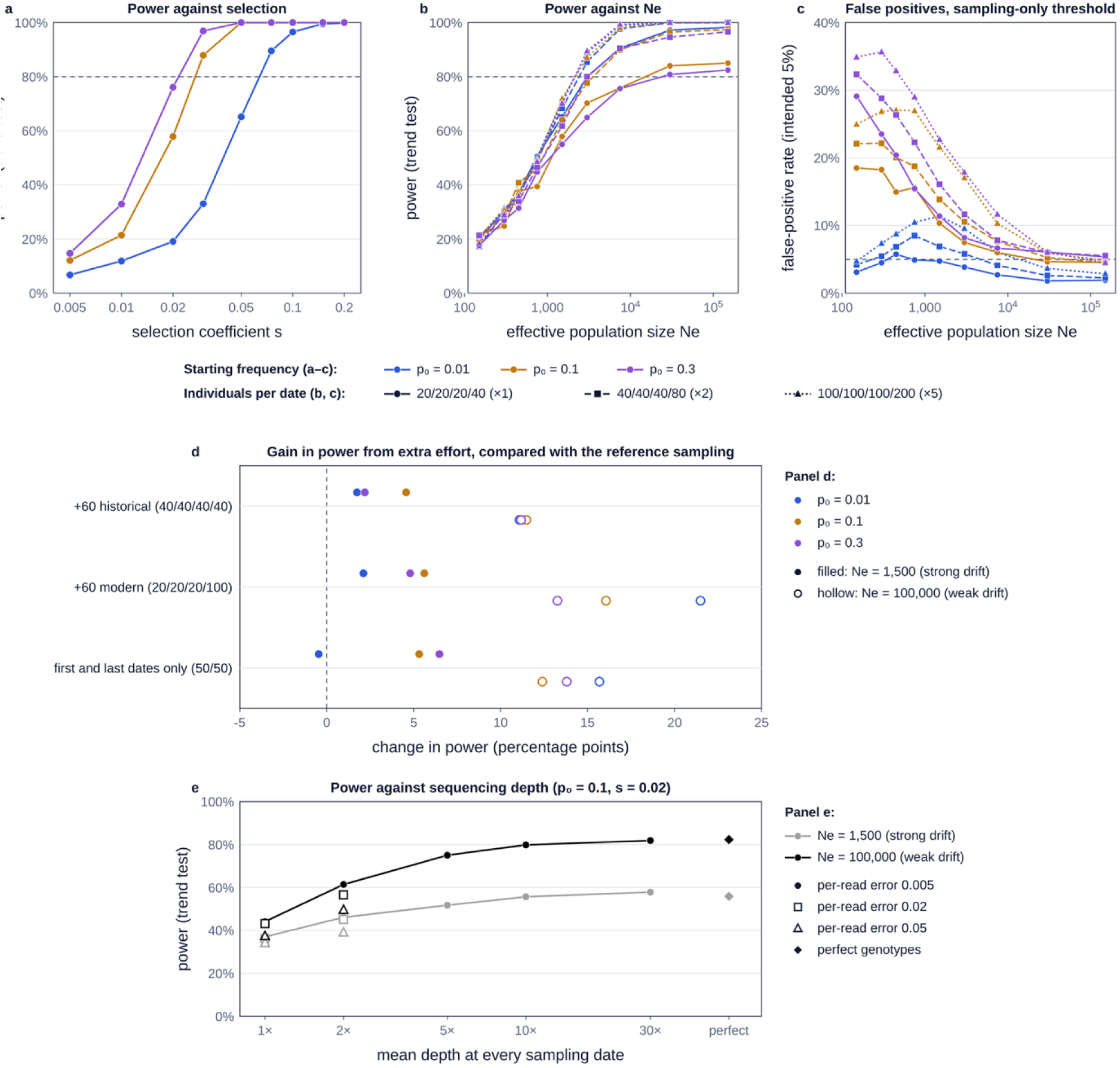
Museum insect case study. Reference design: haplodiploid, *N*_f_=*N*_m_=1,000 (*N*_e_=1,500); 20, 20, 20 and 40 females sampled in 1935, 1955, 1975 and 2025; mean depth 10× for the historical and 15× for the modern samples; trend test with the drift-aware threshold, *α*=0.05. In panels a–d, colours give the starting frequency (*p*_0_=0.01, 0.1 and 0.3). (a) Power against selection coefficient (*N*_e_=1,500). (b) Power and (c) false-positive rate of the sampling-only threshold against *N*_e_, with *s*=0.05, 0.02 and 0.015 for *p*_0_=0.01, 0.1 and 0.3 (as for the filled symbols in d), for the reference samples and for two and five times as many individuals at every date (line style and marker). (d) Change in power relative to the reference sampling from adding 60 historical or 60 modern specimens, or from sampling the same 100 individuals at the first and last dates only; filled symbols, *N*_e_ 1,500; hollow, 100,000; at each *N*_e_, *s* was chosen so that the reference design has 55–67% power. (e) Power against the mean depth at every sampling date (1× to 30×, and perfect genotypes) at *N*_e_=1,500 (grey) and 100,000 (black) (*p*_0_=0.1, *s*=0.02, minimum depth 1 read), with a per-read error rate of 0.005 and, at 1× and 2×, of 0.02 and 0.05 (marker shape). Each point is based on 1,000 simulations under selection in a–c (standard deviation of power about 1.5 percentage points) and 5,000 in d and e; thresholds in e use 10,000 neutral simulations.

Drift limited what additional sampling could achieve (Figure 3b). With strong drift (*N*_e_=150), power was 18–22% for the three starting frequencies, and five times as many individuals at every date changed it by less than 1 point. At the same *N*_e_, the sampling-only threshold gave more false positives with more individuals: for *p*_0_=0.1, 18.5% with the reference samples and 24.9% with five times as many, and for *p*_0_=0.3, 29.1% and 34.9%; for the rare allele (*p*_0_=0.01), it stayed below 5% (3.1% and 4.7%; Figure 3c). With weak drift, in contrast, additional effort increased power more, and the increase depended on where the effort was added (Figure 3d; Table S9). At *N*_e_=100,000, adding 60 historical specimens increased power by 11.1–11.5 points and adding 60 modern specimens by 13.3–21.5, compared with 1.7–4.6 and 2.1–5.6 at 1,500. Concentrating the same 100 individuals at the first and last dates increased power by 12.4–15.7 points at 100,000, compared with −0.5 to +6.5 at 1,500.

Sequencing the whole design at very low depth reduced power, much more so when drift was weak (Figure 3e). With *s*=0.02 and a minimum depth of 1, power at *N*_e_=1,500 was 58% at 30×, 56% at 10×, 52% at 5×, 46% at 2× and 37% at 1×; at *N*_e_=100,000, it fell from 82% at 30× to 75% at 5×, 61% at 2× and 44% at 1×. Low depth adds genotyping noise (missing genotypes and miscalled heterozygotes) to sampling error. When drift is weak, sampling and genotyping are the main sources of chance, so this added noise reduces power strongly. When drift is strong, drift is the limiting factor and the added noise matters less. Above 10×, tripling the depth added only about 2 points with either level of drift. A higher per-read error rate lowered power further at low depth: with an error rate of 0.05 instead of 0.005, power at 1× was 34% instead of 37% with strong drift and 37% instead of 44% with weak drift. Low depth also raised the false-positive rate of the sampling-only threshold when drift was weak, from 5.4% at 30× to 11.5% at 1× (*N*_e_=100,000). With the default minimum depth of 3, most individuals had fewer than 3 reads at the locus and received no genotype (92% at 1× and 68% at 2×, leaving about 8 and 32 of the 100 sampled individuals), and power fell to 18% at 1× and to 33–38% at 2×.

#### Human ancient DNA

Deeper sequencing of the ancient samples increased power moderately (Figure S6a): with *s*=0.0075, power rose from 73% at 1× to 84% at 10× (82% with perfect genotypes), and with *s*=0.005 from 44% to 51%. With *s*=0.02 or more, power was 100% at all depths. With pseudo-haploid calls, power was 6–12 points lower and increased less with depth (66% at 1× and 72% at 10× with *s*=0.0075; S7.3). With the same sequencing effort of 100 reads per ancient date (Figure S6b; Table S10), power decreased only slightly from 100 individuals at 1× to 25 at 4× (from 79% to 76% with *s*=0.0075), then faster, to 73% with 20 individuals at 5× and 66% with 10 at 10×.

## 4. Discussion

SPOT estimates the power of a temporal study to detect selection before its samples are collected or sequenced, accounting for drift as well as sampling and sequencing. Within its assumptions, it agreed with theoretical expectations (S3) and with SLiM in allele-frequency distributions, power and false-positive rates.

The central element of SPOT is the drift-aware threshold. Because it is set from neutral simulations that include the drift, sampling and genotyping of the design, it gave, on average, close to the intended false-positive rate in all test cases, including strong drift, related individuals and up to 1,000 candidate loci. The sampling-only threshold, which accounts for sampling error but not drift, is known to give more false positives than intended when drift is substantial (Waples, 1989; Wiberg et al., 2017; Spitzer et al., 2020), and our comparison with SLiM shows that this difference is not specific to SPOT. Drift, related individuals and low-depth genotype calls spread the neutral values of Z more widely than the sampling of independent gene copies predicts, so the fixed threshold is too low: more neutral simulations than intended exceed it, and so do more selection simulations, which inflates power. For the reference case with s=0.02, the sampling-only threshold reported a power of 69% instead of 55%. The difference was particularly large when the samples contained chance variation not due to sampling: in small or declining populations, when relatives were sampled together, at low depth and, less intuitively, when more individuals were sampled. Sampling more individuals does not change drift; it reduces sampling error, so drift becomes a larger part of what is observed, and a threshold that ignores it becomes more misleading. Conversely, with weak drift or a rare allele, this threshold was close to its intended rate or below it; for a rare allele, the few copies sampled allow only a few distinct values of Z, so that this fixed threshold cannot reach the intended rate exactly, whereas the drift-aware threshold reaches it by sharing tied values (S1.10). The drift-aware threshold is, however, only as reliable as the scenario it is computed from: if *N*_e_ is overestimated, for example when census numbers are entered as breeders, the neutral simulations contain too little drift, and the threshold would give more false positives than intended.

The case studies show what limits power and which changes to the design improve it. A rare favoured allele is harder to detect than a common one: in the museum insect case study, 80% power required a selection coefficient about three times larger when the allele started at 1% than at 30% (Figure 3a). How much additional sampling helps depends on drift. When drift was strong, five times as many individuals barely increased power (*N*_e_=150; Figure 3b), and adding 60 historical or modern specimens increased it by at most about 6 points (*N*_e_=1,500); when drift was weak, the same 60 specimens increased it by up to 22 points, modern specimens at least as much as historical ones (Figure 3d). Concentrating the same specimens at the first and last dates helped mostly when drift was weak (with selection acting from the first date). Moderate changes in depth mattered less than the number of individuals: above 10×, tripling the depth of all samples added about 2 points (Figure 3e), and for human ancient DNA, with a fixed number of reads per date, power was similar from 100 individuals at 1× to 25 at 4× and fell with fewer, more deeply sequenced individuals (Figure S6b). At the extremes, however, depth mattered, especially when drift was weak: sequencing the whole museum design at 1× instead of 30× reduced power by 21 points with strong drift but by 38 points with weak drift, and damage-like read errors reduced it further (Figure 3e). This extends to the conclusion of Buerkle and Gompert (2013) that, for estimating allele frequencies, many individuals at low coverage are more informative than few at high coverage. Finally, because power depends on the number of generations rather than years, a bacterial pathogen sampled over 17,000 generations reached 80% power with s≈0.0003 (S6). These patterns are expected from theory; SPOT gives their magnitude for a particular design, through the power curve and the sample-size planner (Figure 1c,d).

Like any power analysis, SPOT’s estimates depend on the assumed scenario, and some of its parameters are rarely known precisely, in particular *N*_e_, the starting frequency of the favoured allele and the strength of selection. Users can compute power over a plausible range of each parameter with the power curve (Figure 1c) and choose designs that remain adequate across this range. *N*_e_ can be narrowed with existing information, such as published estimates, temporal estimates from earlier genetic data (Waples, 1989), or the loss of genetic diversity observed in genome-wide data from earlier work or a pilot study, which users can match with the loss reported by SPOT. Finally, each power estimate varies between runs, with a standard deviation of about 1.5–2.2 points with the default 1,000 selection simulations (Section 3.3; S5). Designs whose powers differ by only a few points should therefore be compared with more selection simulations, for example 5,000, especially for very long series, in which single runs varied by more than ten points (S6).

SPOT is not meant to replace general simulators such as SLiM. It is designed to be easy and fast to use, through a graphical interface, for a wide range of temporal designs, at the cost of a deliberately simple model: one locus under constant directional selection from a chosen date, in a single population with discrete, non-overlapping generations, with Poisson-distributed read depths and one per-read error rate for the historical dates and another for the last date, the same for both alleles, analysed with a trend test on called genotypes. Post-mortem damage can create a spurious trend; SPOT assumes it has been removed, for example by excluding transitions. Linked selection, population structure, admixture, overlapping generations or genome-wide scans require dedicated simulations, for example with SLiM. Power also depends on the method that will be used to analyse the data: likelihood-based methods that estimate *s* from the whole allele-frequency trajectory (Mathieson & McVean, 2013; Schraiber et al., 2016), or that keep the uncertainty of each genotype (genotype likelihoods; Korneliussen et al., 2014) instead of calling a single genotype per individual as SPOT does, may have different power for the same design. With pseudo-haploid calls, power is lower, because each individual then gives one gene copy instead of two (S7.3).

Within this scope, SPOT answers in seconds, and without programming, a question that often remains open until the data have been collected: can this design detect selection of plausible strength? Answering it before collecting or sequencing samples, or before adding to an existing collection, allows researchers to compare alternative designs and to spend scarce specimens and sequencing effort where they increase power most.

## Author Contributions

Y.B.C. and I.B. conceived the study. Y.B.C. developed the software, ran the simulations and analyses, and wrote the manuscript. I.J., G.B., A.P., B.P., and I.B. tested the software. I.B., I.J., B.P., and G.B. acquired the funding. I.J., G.B., A.P., B.P., I.B., and Y.B.C. revised the manuscript and approved the final version.

## Acknowledgements

We thank Victoria Mullin and William Marsh for testing SPOT and providing suggestions for its improvement.

## Ethics Statement

The authors have nothing to report.

## Data Availability Statement

*Data and code.* SPOT is available at https://yacinebenchehida.github.io/SPOT-simulator/. Its source code, documentation, and the scripts and results of all analyses presented in this article (with commands provided in S8) are available at https://github.com/yacinebenchehida/SPOT-simulator (MIT licence). SPOT is archived on Zenodo (https://doi.org/10.5281/zenodo.24423881). This study used only simulated data.

*Benefit-Sharing Statement.* Not applicable: no genetic resources or samples were used. SPOT is free, open-source software available to all researchers.

## Use of generative AI

The large language model assistant Claude (Anthropic; Opus 5.5) was used during this work, under the supervision and direction of the authors, to: (i) revise parts of the code of the SPOT application and the validation scripts; (ii) verify the commands used in the simulation analyses; and (iii) draft and edit parts of the manuscript and user documentation. All decisions regarding the study design, the model, the analyses and their interpretation were made by the authors. The authors reviewed all AI-assisted code and text, and take full responsibility for the content of this publication.

## References

Anchieri, L., Amorim, C. E. G., Neuenschwander, S., & Malaspinas, A.-S. (2026). Assessing ancient DNA sampling strategies for natural selection inference in humans using allele frequency time series data. Genome Biology and Evolution, 18(2), evaf234. 10.1093/gbe/evaf234

Armitage, P. (1955). Tests for linear trends in proportions and frequencies. Biometrics, 11(3), 375–386. 10.2307/3001775

Barata, C., Borges, R., & Kosiol, C. (2023). Bait-ER: A Bayesian method to detect targets of selection in Evolve-and-Resequence experiments. Journal of Evolutionary Biology, 36(1), 29–44. 10.1111/jeb.14134

Briggs, A. W., Stenzel, U., Johnson, P. L. F., Green, R. E., Kelso, J., Prüfer, K., Meyer, M., Krause, J., Ronan, M. T., Lachmann, M., & Pääbo, S. (2007). Patterns of damage in genomic DNA sequences from a Neandertal. Proceedings of the National Academy of Sciences, 104(37), 14616–14621. 10.1073/pnas.0704665104

Buerkle, C. A., & Gompert, Z. (2013). Population genomics based on low coverage sequencing: How low should we go? Molecular Ecology, 22(11), 3028–3035. 10.1111/mec.12105

Cheng, X., & Steinrücken, M. (2025). diplo-locus: A lightweight toolkit for inference and simulation of time-series genetic data under general diploid selection. G3: Genes, Genomes, Genetics, 15(8), jkaf123. 10.1093/g3journal/jkaf123

Clark, R. D., Catalano, K. A., Fitz, K. S., Garcia, E., Jaynes, K. E., Reid, B. N., Sawkins, A., Snead, A. A., Whalen, J. C., & Pinsky, M. L. (2025). The practice and promise of temporal genomics for measuring evolutionary responses to global change. Molecular Ecology Resources, 25(5), e13789. 10.1111/1755-0998.13789

Cochran, W. G. (1954). Some methods for strengthening the common χ² tests. Biometrics, 10(4), 417–451. 10.2307/3001616

Dehasque, M., Ávila-Arcos, M. C., Díez-del-Molino, D., Fumagalli, M., Guschanski, K., Lorenzen, E. D., Malaspinas, A.-S., Marques-Bonet, T., Martin, M. D., Murray, G. G. R., Papadopulos, A. S. T., Therkildsen, N. O., Wegmann, D., Dalén, L., & Foote, A. D. (2020). Inference of natural selection from ancient DNA. Evolution Letters, 4(2), 94–108. 10.1002/evl3.165

Dunn, O. J. (1961). Multiple comparisons among means. Journal of the American Statistical Association, 56(293), 52–64. 10.1080/01621459.1961.10482090

Feder, A. F., Kryazhimskiy, S., & Plotkin, J. B. (2014). Identifying signatures of selection in genetic time series. Genetics, 196(2), 509–522. 10.1534/genetics.113.158220

Ferrer-Admetlla, A., Leuenberger, C., Jensen, J. D., & Wegmann, D. (2016). An approximate Markov model for the Wright–Fisher diffusion and its application to time series data. Genetics, 203(2), 831–846. 10.1534/genetics.115.184598

Foll, M., Shim, H., & Jensen, J. D. (2015). WFABC: A Wright–Fisher ABC-based approach for inferring effective population sizes and selection coefficients from time-sampled data. Molecular Ecology Resources, 15(1), 87–98. 10.1111/1755-0998.12280

Good, B. H., McDonald, M. J., Barrick, J. E., Lenski, R. E., & Desai, M. M. (2017). The dynamics of molecular evolution over 60,000 generations. Nature, 551(7678), 45–50. 10.1038/nature24287

Haller, B. C., Ralph, P. L., & Messer, P. W. (2026). SLiM 5: Eco-evolutionary simulations across multiple chromosomes and full genomes. Molecular Biology and Evolution, 43(1), msaf313. 10.1093/molbev/msaf313

He, Z., Dai, X., Lyu, W., Beaumont, M., & Yu, F. (2023). Estimating temporally variable selection intensity from ancient DNA data. Molecular Biology and Evolution, 40(3), msad008. 10.1093/molbev/msad008

Hedrick, P. W., & Parker, J. D. (1997). Evolutionary genetics and genetic variation of haplodiploids and X-linked genes. Annual Review of Ecology and Systematics, 28(1), 55–83. 10.1146/annurev.ecolsys.28.1.55

Hoggart, C. J., Chadeau-Hyam, M., Clark, T. G., Lampariello, R., Whittaker, J. C., De Iorio, M., & Balding, D. J. (2007). Sequence-level population simulations over large genomic regions. Genetics, 177(3), 1725–1731. 10.1534/genetics.106.069088

Kofler, R., & Schlötterer, C. (2014). A guide for the design of evolve and resequencing studies. Molecular Biology and Evolution, 31(2), 474–483. 10.1093/molbev/mst221

Korneliussen, T. S., Albrechtsen, A., & Nielsen, R. (2014). ANGSD: Analysis of next generation sequencing data. BMC Bioinformatics, 15(1), 356. 10.1186/s12859-014-0356-4

Mallick, S., Micco, A., Mah, M., Ringbauer, H., Lazaridis, I., Olalde, I., Patterson, N., & Reich, D. (2024). The Allen Ancient DNA Resource (AADR) a curated compendium of ancient human genomes. Scientific Data, 11(1), 182. 10.1038/s41597-024-03031-7

Mathieson, I., & McVean, G. (2013). Estimating selection coefficients in spatially structured populations from time series data of allele frequencies. Genetics, 193(3), 973–984. 10.1534/genetics.112.147611

Nielsen, R., Paul, J. S., Albrechtsen, A., & Song, Y. S. (2011). Genotype and SNP calling from next-generation sequencing data. Nature Reviews Genetics, 12(6), 443–451. 10.1038/nrg2986

Peng, B., & Kimmel, M. (2005). simuPOP: A forward-time population genetics simulation environment. Bioinformatics, 21(18), 3686–3687. 10.1093/bioinformatics/bti584

Percival-Alwyn, L., Barnes, I., Clark, M. D., Cockram, J., Coffey, M. P., Jones, S., Kersey, P. J., Kidner, C. A., Kosiol, C., Li, B., Marsh, W. A., Zhou, J., Caccamo, M., & Milne, I. (2025). UKCropDiversity-HPC: A collaborative high-performance computing resource approach for sustainable agriculture and biodiversity conservation. Plants, People, Planet, 7(4), 969–977. 10.1002/ppp3.10607

Pinsky, M. L., Eikeset, A. M., Helmerson, C., Bradbury, I. R., Bentzen, P., Morris, C., Gondek-Wyrozemska, A. T., Baalsrud, H. T., Brieuc, M. S. O., Kjesbu, O. S., Godiksen, J. A., Barth, J. M. I., Matschiner, M., Stenseth, N. C., Jakobsen, K. S., Jentoft, S., & Star, B. (2021). Genomic stability through time despite decades of exploitation in cod on both sides of the Atlantic. Proceedings of the National Academy of Sciences, 118(15), e2025453118. 10.1073/pnas.2025453118

Santander, C. G., & Moltke, I. (2025). SimOutbreakSelection: A simulation-based tool to optimise sampling design and analysis strategies for detecting epidemic-driven selection. Nature Communications, 16(1), 6814. 10.1038/s41467-025-61574-8

Schraiber, J. G., Evans, S. N., & Slatkin, M. (2016). Bayesian inference of natural selection from allele frequency time series. Genetics, 203(1), 493–511. 10.1534/genetics.116.187278

Spitzer, K., Pelizzola, M., & Futschik, A. (2020). Modifying the Chi-square and the CMH test for population genetic inference: Adapting to overdispersion. The Annals of Applied Statistics, 14(1), 202–220. 10.1214/19-AOAS1301

Uricchio, L. H., & Hernandez, R. D. (2014). Robust forward simulations of recurrent hitchhiking. Genetics, 197(1), 221–236. 10.1534/genetics.113.156935

Vlachos, C., & Kofler, R. (2018). MimicrEE2: Genome-wide forward simulations of evolve and resequencing studies. PLOS Computational Biology, 14(8), e1006413. 10.1371/journal.pcbi.1006413

Waples, R. S. (1989). A generalized approach for estimating effective population size from temporal changes in allele frequency. Genetics, 121(2), 379–391. 10.1093/genetics/121.2.379

Wiberg, R. A. W., Gaggiotti, O. E., Morrissey, M. B., & Ritchie, M. G. (2017). Identifying consistent allele frequency differences in studies of stratified populations. Methods in Ecology and Evolution, 8(12), 1899–1909. 10.1111/2041-210X.12810

Wright, S. (1931). Evolution in Mendelian populations. Genetics, 16(2), 97–159. 10.1093/genetics/16.2.97

