## Supplementary Information for "Power analysis for selection in allele-frequency time series"

Contents

**S1 | The model simulated by SPOT 3**
S1.1 | Time, starting point and onset of selection 3
S1.2 | One generation, step by step 3
S1.3 | Reproduction and selection 4
S1.4 | Drift 5
S1.5 | Changes in Ne over time 6
S1.6 | Migration 7
S1.7 | Sampling of individuals 7
S1.8 | Sequencing and genotype calls 8
S1.9 | The score Z 10
S1.10 | The threshold and simulations with the same value of Z 11
S1.11 | The sampling-only threshold 13
S1.12 | Number of neutral simulations, and many loci 14
S1.13 | Other outputs 15
**S2 | False-positive rate of the drift-aware threshold 16**
S2.1 | Procedure 16
S2.2 | Test cases 17
S2.3 | Many loci 18
**S3 | Agreement with theoretical expectations 21**
S3.1 | Selection 21
S3.2 | Power without drift 22
S3.3 | Drift: loss of genetic diversity 23
S3.4 | Sampling of relatives 24
**S4 | Comparison with SLiM 25**
S4.1 | SLiM model 26
S4.2 | Scenarios and comparisons 27
S4.3 | Results 28
**S5 | Precision and speed 31**
S5.1 | Procedure 32
S5.2 | Results 33
S5.3 | Practical consequences 33
**S6 | Rescaling of long time series 36**
S6.1 | Rescaling 36
S6.2 | Validation against simulations of every generation 37
S6.3 | Limits of rescaling 39

S6.4 | Example: a bacterial pathogen 40
**S7 | Case studies: tables and figures 41**
S7.1 | Museum insect 41
S7.2 | Human ancient DNA 42
S7.3 | Pseudo-haploid calls 43
**S8 | Reproducibility 44**
**References 45**

### S1 | The model simulated by SPOT

This section describes, step by step, what happens in one simulation. It completes Sections 2.2–2.5 of the main text. Settings of the application are given in quotation marks, with the name they have in the interface. Throughout, "the favoured allele" is the allele that selection makes more frequent, and all frequencies are frequencies of this allele.

#### *S1.1 | Time, starting point and onset of selection*

SPOT counts time in generations from the first sampling date, which is generation 0. The generation of each later sampling date is the number of years since the first date divided by the generation time ("Years per generation"), rounded to the nearest whole number. For example, in the reference case (generation time 1 year), the sampling years 1935, 1955, 1975 and 2025 are generations 0, 20, 40 and 90; in the human ancient DNA case study (28 years per generation), 7000 BCE and 2000 CE are 321 generations apart.

A simulation starts at the first sampling date: the favoured allele then has the frequency set in "Allele frequency at the first point ($p_0$)", written $p_0$, in breeding females and in breeding males (in haploids, which have no sexes, in all breeding individuals). Nothing before the first sampling date is simulated; any earlier history of the population is summarised by $p_0$.

Selection acts in the generations that come after the year set in "Selection starts in year". Three situations are possible. (i) The onset is the first sampling date (the default): selection acts from generation 1, so the first sample, taken in generation 0, is taken before any selection, and every later sample reflects selection and drift. (ii) The onset is earlier than the first sampling date: because the simulation starts at the first date, this is equivalent to (i); the effect of selection before the first date is contained in $p_0$. (iii) The onset is later than the first sampling date: the population then changes by drift only until the onset, and by selection and drift afterwards. For example, in the reference case with selection starting in 1975, generations 1–40 are neutral and selection acts from generation 41, so the first three samples contain no effect of selection.

#### *S1.2 | One generation, step by step*

Each generation goes through five steps, in this order, described in the sections below:

(1) Reproduction (Section S1.3). The breeding adults of the generation produce offspring.

(2) Selection (Section S1.3). If selection acts in this generation, the offspring survive according to their genotype.

(3) Sampling and genotyping (Sections S1.7 and S1.8). If the generation is a sampling date, the sampled individuals are drawn from the offspring that survived selection, and their genotypes are called from simulated sequencing reads.

(4) Drift (Sections S1.4 and S1.5). The breeding adults of the next generation are drawn at random, in finite numbers, from the offspring that survived selection.

(5) Migration (Section S1.6). A fraction of these breeding adults is replaced by migrants.

#### *S1.3 | Reproduction and selection*

SPOT follows the frequency of the favoured allele among breeding females and among breeding males, written $p_f$ and $p_m$. It does not follow individual organisms: the genotype frequencies of the offspring are calculated from $p_f$ and $p_m$, assuming random mating.

*Selection.* Selection acts on the survival of offspring (viability selection). Its strength is set by the selection coefficient $s$ ("Selection coefficient s"), the relative survival advantage of individuals that carry the favoured allele. In diploid individuals, the dominance coefficient $h$ ("Dominance in diploid individuals") sets the advantage of heterozygotes: individuals with zero, one and two copies of the favoured allele survive in proportions 1, $1 + hs$ and $1 + s$. With $h = 0$ (recessive), only homozygotes for the favoured allele have an advantage; with $h = 0.5$ (additive), heterozygotes are exactly intermediate; with $h = 1$ (dominant), one copy gives the full advantage. Haploid individuals survive in proportions 1 without the favoured allele and $1 + s$ with it. After selection, the frequency of each genotype is its frequency before selection multiplied by its survival, divided by the average survival of all offspring, so that the frequencies again sum to 1. In males, $s$ is multiplied by the setting "Selection in males (× s)" (1 by default, that is, the same selection in both sexes); this selection coefficient in males is written $s_m$.

*Diploids.* Each offspring receives one allele from its mother and one from its father. Before selection, offspring therefore have two copies of the favoured allele with frequency $P_2 = p_f p_m$, one copy with frequency $P_1 = p_f(1 - p_m) + p_m(1 - p_f)$, and no copy with frequency $P_0 = (1 - p_f)(1 - p_m)$. After selection, each genotype is weighted by its survival ($1 + s$, $1 + hs$ and 1; see Selection, above), and the frequency of the favoured allele among the offspring becomes

$$p' = \frac{P_2(1+s)+\frac{1}{2}P_1(1+hs)}{P_2(1+s)+P_1(1+hs)+P_0} \quad (1)$$

where the numerator counts the copies of the favoured allele carried by the surviving offspring (two per offspring with two copies, one per heterozygote, hence the factor ½) and the denominator is the average survival of all offspring. Selection is then applied separately to female offspring (with $s$) and to male offspring (with $s_m$).

*Haplodiploids.* Daughters are diploid, as above. Sons carry only their mother's allele, so the frequency of the favoured allele among sons is $p_f$ before selection, and after selection it is given by Equation (2), with $p_f$ in place of $p$ and $s_m$ in place of $s$.

*Haploids.* Each individual carries one allele and reproduces clonally, and there are no sexes. With frequency $p$ before selection, individuals with the favoured allele (a proportion $p$) survive in proportion $1 + s$ and the others in proportion 1, so the average survival is $1 + ps$ and the frequency after selection is

$$p' = \frac{p(1+s)}{1+ps} \quad (2)$$

#### *S1.4 | Drift*

Drift arises because only a finite number of offspring become the breeding adults of the next generation. SPOT draws these breeding adults at random from the offspring that survived selection. The numbers of breeding females and males are set by "Breeding females per generation (Nf)" and "Breeding males per generation (Nm)", written $N_f$ and $N_m$ (in haploids, "Breeding individuals per generation (N)", written $N$). They are numbers of breeding adults, not census sizes: together they determine the *effective population size ($N_e$; Section 2.2 of the main text)*.

For breeding females, SPOT draws the number of their $2N_f$ gene copies that carry the favoured allele from a binomial distribution, with $2N_f$ draws and a probability equal to the frequency of the favoured allele in female offspring after selection; the new $p_f$ is this number divided by $2N_f$. Breeding males are drawn in the same way from male offspring, with $2N_m$ gene copies in diploids and $N_m$ in haplodiploids (each male carries one copy). In haploids, $N$ gene copies are drawn. With "No drift (infinite population)", this step is skipped, and allele frequencies change by selection and migration only.

*A faster draw for large populations.* Each of these draws takes a number $n$ of gene copies, each carrying the favoured allele with probability $q$, the frequency of the favoured allele among the offspring after selection. The number drawn follows a binomial distribution with mean $nq$ and variance $nq(1 - q)$. Drawing it exactly takes longer as $n$ grows. When at least 20 copies of the rarer allele are expected, that is, when $n \times \min(q, 1 - q) \geq 20$, SPOT therefore draws the number from a normal distribution with the same mean and variance, and rounds it to the nearest whole number. When $n$ is large and $q$ is not too close to 0 or 1, the binomial distribution is close to a normal distribution (de Moivre–Laplace theorem). The usual criterion is that both alleles are expected in large numbers, which the threshold of 20 copies of the rarer allele ensures. For example, the 1,000 breeding females of a diploid population carry n = 2,000 gene copies. If the frequency of the favoured allele among female offspring after selection is q = 0.01, 2,000 × 0.01 = 20 copies of the favoured allele are expected, and the number drawn comes from the normal distribution; the same holds for any q between 0.01 and 0.99. If q = 0.005, only 2,000 × 0.005 = 10 copies are expected (fewer than 20 copies of the rarer allele), and the number is then drawn exactly from the binomial distribution. The same applies at q = 0.995, where the other allele is the rarer one: 1,990 copies of the favoured allele are expected, but only 10 of the other allele.

#### ***S1.5 | Changes in*** $N_e$ **over time**

By default, the numbers of breeding adults are the same in every generation. Two settings make them change over time; in both, the numbers of breeding females and males of each generation are rounded to whole numbers and are never below 2.

*Gradual change.* "Ne at the last point relative to the first" sets the ratio of the breeding numbers, and therefore of $N_e$, at the last sampling date to those at the first (1 by default). The breeding numbers then change exponentially between the first and last sampling dates. Writing $R$ for this ratio and $G$ for the generation of the last sampling date, the numbers of breeding females and males in generation $g$ are $N_f \times R^{g/G}$ and $N_m \times R^{g/G}$: they equal $N_f$ and $N_m$ at the first sampling date ($g = 0$), $R$ times these numbers at the last ($g = G$).

*A chosen* $N_e$ *in each interval.* With "Set Ne separately for each interval", *the user gives a value of* $N_e$ *for each interval* between two consecutive sampling dates, for example to represent a bottleneck. In each interval, SPOT multiplies $N_f$ and $N_m$ by the same factor, $N_{e,j}/N_e$, where $N_{e,j}$ *is the value of* $N_e$ *chosen for interval j* and $N_e$ *the value given by* $N_f$ and $N_m$. $N_e$

*in the interval then equals* the chosen value, and the ratio of females to males is kept. *The value of* $N_e$ *given by* $N_f$ and $N_m$ is $4N_fN_m/(N_f + N_m)$ in diploids, $9N_fN_m/(4N_m + 2N_f)$ in haplodiploids (Hedrick & Parker, 1997) and $N$ in haploids.

#### *S1.6 | Migration*

After drift, a fraction of the breeding adults, set by "Migrants per generation, m" and written $m$, is replaced by migrants from a source population in which the favoured allele has a fixed frequency, set by "Allele frequency in the migrant source" and written $p_{mig}$ (by default $p_{mig} = p_0$, that is, the allele is not selected in the source). The frequency of the favoured allele among breeding females then becomes $(1 - m)p_f + m\,p_{mig}$, and that among breeding males $(1 - m)p_m + m\,p_{mig}$: a fraction $1 - m$ of the breeding adults keeps the frequency of the population, and a fraction $m$ has the frequency of the source.

#### *S1.7 | Sampling of individuals*

At each sampling date, SPOT samples the number of individuals set in the sampling design, from the offspring that survived selection in that generation. Because SPOT follows allele frequencies rather than individual organisms, the sampled individuals do not exist before they are sampled: SPOT creates them at the sampling date, drawing their genotypes from the genotype frequencies of the population at that time. For families, the two parents are created in the same way, only to generate the siblings. These individuals are used only to produce the sample: they do not take part in reproduction or drift, and do not change the allele frequencies of the population.

*Sex of sampled individuals.* In diploids and haplodiploids, each sampled individual is a male with probability set by "Males in the sample" (0 by default, that is, females only), and a female otherwise. In haplodiploids, sampled males carry a single allele. In diploid families (see Relatives, below), the sex of siblings is not drawn: siblings are not subject to selection, so their sex has no effect. Haploids have no sexes.

*Unrelated individuals.* With "Sampled individuals per family" set to 1 (the default), the sampled individuals are unrelated: each individual's genotype is drawn independently from the genotype frequencies, after selection, of the offspring of its sex.

*Relatives.* With "Sampled individuals per family" set to a number larger than 1, individuals are sampled in families of full siblings, which share both parents. The setting gives the number of siblings in each family. For example, with 20 individuals at a date, a setting of 2 gives 10 pairs of siblings, and a setting of 4 gives 5 families of 4 siblings. When the number of individuals is not a multiple of the setting, the last family of the date has fewer siblings: with 20 individuals and a setting of 3, the date has 6 families of 3 siblings and 1 family of 2. Each family is generated as follows. (1) The mother's genotype is drawn from the genotype frequencies of female offspring after selection, and the father's from those of male offspring (a haplodiploid father carries one allele). (2) Each sibling receives one allele from its mother and one from its father, following Mendel's rules: a parent with two copies of the favoured allele always transmits it, a parent with none never does, and a heterozygous parent transmits it with probability 0.5. A haplodiploid father transmits his single allele to all his daughters, and haplodiploid sons receive their mother's allele only. (3) The siblings are not subject to further selection. Because siblings share their parents' alleles, a family of 4 siblings carries less information on the allele frequency of the population than 4 unrelated individuals. In haploids, this setting has no effect.

#### *S1.8 | Sequencing and genotype calls*

In real studies, genotypes are not observed directly but inferred from sequencing reads, and at low depth they can be wrong: a heterozygote whose reads all happen to come from the same chromosome copy looks homozygous. SPOT simulates this process so that its effect on power is included. With "Perfect genotypes (ignore sequencing)", this step is skipped and every sampled individual has its true genotype.

Otherwise, SPOT simulates the sequencing of each sampled individual in three steps. (1) *Number of reads.* The individual receives a number of reads covering the locus, written $d$. In real data, the number of reads varies between individuals sequenced at the same mean depth, so $d$ is drawn from a Poisson distribution whose mean is the mean depth of the sampling date: "Mean depth, earlier points" for all dates except the last, and "Mean depth, last point" for the last. For example, at a mean depth of 10×, one individual may receive 7 reads and another 13. When the mean exceeds 60, a normal distribution with the same mean and variance, rounded to a whole number, is used instead, because it is faster and practically identical (Section S1.4). (2) *Minimum depth.* If $d$ is below "Minimum depth to call a genotype", the individual receives no genotype and is left out of the counts of its sampling date, as it would be in a real study. (3) *Sequencing errors.* Sequencing is not perfect: a read can show an allele that the individual does not carry. SPOT simulates these errors with a per-read error rate, written $\varepsilon$, the probability that a read shows the wrong allele (for example, one read in 200 when $\varepsilon = 0.005$), set separately for the earlier dates and the last date ("Per-read error rate, earlier points" and "Per-read error rate, last point") because historical samples often carry more errors; because the locus has two alleles in diploids, an error always shows the other allele. For each read of an individual, the probability that the read shows the favoured allele, written $\rho$, therefore depends on the genotype of this individual at the locus. If the individual is homozygous for the favoured allele (it carries two copies of it), every read comes from a copy that carries the favoured allele, so each read shows the favoured allele unless it has an error: $\rho = 1 - \varepsilon$.

If the individual is heterozygous (it carries one copy of the favoured allele and one copy of the other allele), a read can show the favoured allele in two ways. Either the read comes from the copy that carries the favoured allele (probability 0.5) and has no error (probability $1 - \varepsilon$), which gives $0.5(1 - \varepsilon)$; or the read comes from the copy that carries the other allele (probability 0.5) and has an error, so that it shows the favoured allele (probability $\varepsilon$), which gives $0.5\varepsilon$. Adding the two: $\rho = 0.5(1 - \varepsilon) + 0.5\varepsilon = 0.5$. If the individual is homozygous for the other allele (it carries no copy of the favoured allele), the read shows the favoured allele only if it has an error: $\rho = \varepsilon$. Reads are independent of each other, so for an individual with $d$ reads, the number of reads that show the favoured allele, written $r$, is drawn from a binomial distribution with $d$ trials and probability $\rho$.

From these reads, SPOT then calls the genotype of the individual, that is, it decides which genotype the individual has, as a genotype caller would in a real study. It calls the genotype under which the observed reads are most probable (maximum-likelihood calling; Nielsen et al., 2011). For a given set of $d$ reads of which $r$ show the favoured allele, this probability is $\varepsilon^r(1 - \varepsilon)^{d - r}$ for a homozygote with no copy (the $r$ reads are errors and the $d - r$ others are correct), $0.5^d$ for a heterozygote (each read shows either allele with probability 0.5), and $(1 - \varepsilon)^r\varepsilon^{d - r}$ for a homozygote with two copies (the $r$ reads are correct and the $d - r$ others are errors). If the heterozygote and a homozygote are equally probable, the heterozygote is called.

*Calling haploid individuals.* For haploids and haplodiploid males, the allele shown by most reads is called; if both alleles are shown by the same number of reads, one of them is chosen at random.

*Pseudo-haploid calls.* With "Genotype calls" set to "Pseudo-haploid", no genotype is called. For each individual with at least the minimum depth, one of its $d$ reads is drawn at random, and the allele it shows counts as one gene copy, in diploid as in haploid individuals. This is the same as drawing a single read, which shows the favoured allele with probability $\rho$ as above, so additional reads matter only by bringing more individuals above the minimum depth. As usual, the drift-aware threshold is set from neutral simulations analysed in the same way. For diploid individuals, the first two summaries described below are then not shown; the proportion of individuals below the minimum depth still is. Section S7.3 compares power with both kinds of calls in the human ancient DNA case study.

SPOT reports three summaries of this step in the interface: the proportion of true heterozygotes among the sampled individuals that received a genotype and were called heterozygous ("Heterozygous individuals called heterozygous"; for haploid individuals, the proportion of gene copies called correctly), the same proportion for true homozygotes ("homozygous individuals called correctly"), and the proportion of sampled individuals left without a genotype because their depth is below the minimum ("of sampled individuals below the minimum depth"). All three are averaged over the selection simulations of the run.

#### ***S1.9 | The score Z***

For each simulated data set, SPOT summarises the called genotypes by the score $Z$ of the Cochran–Armitage trend test (Cochran, 1954; Armitage, 1955), a standard test of whether a proportion changes linearly with time. Let $K$ be the number of sampling dates and, for sampling date $i$ ($i = 1, \ldots, K$):

$a_i$, the number of gene copies with a called genotype: two for each called diploid individual and one for each called haploid individual (a haploid or a haplodiploid male), or one per individual with pseudo-haploid calls (Section S1.8);

$x_i$, the number of these gene copies that carry the favoured allele, so that $x_i/a_i$ is the frequency of the favoured allele in the sample;

$t_i$, the time of the sampling date, in generations since the first sampling date (Section S1.1).

Then

$$Z = \frac{\sum_i x_i (t_i - \bar{t})}{\sqrt{\bar{p}(1-\bar{p}) \sum_i a_i (t_i - \bar{t})^2}} \quad (3)$$

where the sums run over the sampling dates, $\bar{p} = \Sigma_i x_i / \Sigma_i a_i$ is the frequency of the favoured allele in all samples pooled, and $\bar{t} = \Sigma_i a_i t_i / \Sigma_i a_i$ is the average sampling time, in which each date counts in proportion to its number of called gene copies. The numerator measures how much the favoured allele is more frequent in later samples than in earlier ones; the denominator is its expected standard deviation if the sampled gene copies were independent and the frequency did not change. $Z$ is therefore positive when the favoured allele rises, and larger when the rise is larger or when more gene copies are sampled.

SPOT computes $Z$ twice: with all sampling dates (the trend test) and with the first and last sampling dates only (the first-versus-last test).

$Z$ cannot be computed when no gene copy is called (which happens after genotype calling, when all sampled individuals have fewer reads than the minimum depth; Section S1.8), or when the favoured allele is absent from all samples or present in all called gene copies ($\bar{p} = 0$ or 1), because the denominator is then zero. In these cases, SPOT sets $Z$ to 0, that is, no evidence of a rise. This happens often for a rare allele: with $p_0 = 0.01$, a large proportion of simulations have no copy of the favoured allele in any sample, and all of them have $Z = 0$.

#### *S1.10 | The threshold and simulations with the same value of Z*

*The threshold.* SPOT sets the threshold of each test from its neutral simulations (Section 2.1 of the main text). With 2,000 neutral simulations and a significance level of 5%, the threshold is placed so that 100 of the 2,000 neutral simulations (5%) are above it; more generally, a proportion $\alpha'$ of the $n$ neutral simulations, that is, $\alpha' n$ simulations, where $\alpha'$ is the significance level per locus. In practice, the $n$ neutral values of $Z$ are sorted from smallest to largest, and the threshold is the value at position $(1 - \alpha')n$, rounded up to a whole number. A simulation, neutral or with selection, is declared a detection when its $Z$ is above the threshold.

*The problem: simulations with the same value of Z.* $Z$ is calculated from counts of gene copies, so it can only take a limited number of values. When many gene copies are sampled at several dates, these values are so numerous that two neutral simulations rarely have exactly the same $Z$, and the threshold can be placed with exactly $\alpha' n$ neutral simulations above it.

When few copies of the favoured allele are sampled (for a rare allele, or with few individuals) or when $Z$ is computed from two dates only (the first-versus-last test), many simulations share exactly the same value of $Z$, and the threshold may fall on such a value. No threshold can then separate the simulations that share this value, and the proportion of neutral simulations counted as detections cannot be exactly $\alpha'$: counting only the simulations strictly above the threshold gives fewer than $\alpha' n$ detections, so the test gives fewer false positives than intended (it is conservative) and underestimates power; counting the tied simulations as well gives more than $\alpha' n$ detections.

*The rule used by SPOT.* Let $A$ be the number of neutral simulations whose $Z$ is above the threshold, and $B$ the number whose $Z$ is exactly equal to it (the threshold is itself one of the neutral values, so $B$ is at least 1). The threshold is chosen so that $A \leq \alpha' n \leq A + B$. SPOT counts each simulation whose $Z$ is above the threshold as one detection, and each simulation whose $Z$ equals the threshold as a fraction $\gamma$ of a detection, with

$$\gamma = \frac{\alpha' n - A}{B} \tag{4}$$

a number between 0 and 1. The neutral simulations then give $A + \gamma B = \alpha' n$ detections, that is, a false-positive rate of exactly $\alpha'$. The rule does not move the threshold: it only sets how the simulations whose $Z$ equals the threshold are counted. The same rule, with the same $\gamma$, is applied to the selection simulations, because whether a simulation counts as a detection depends only on its value of $Z$, not on whether it was simulated with or without selection; the false-positive rate and the power then describe the same test. Power is $(S_> + \gamma S_=)/R$, where $S_>$ and $S_=$ are the numbers of selection simulations whose $Z$ is above and equal to the threshold, and $R$ is the number of selection simulations. When no other neutral simulation has the same $Z$ as the threshold ($B = 1$ and $A = \alpha' n$), $\gamma = 0$ and the rule changes nothing.

*Example.* In one run of the example without drift (diploid population without drift, $p_0 = 0.01$, unrelated individuals, perfect genotypes; Section S1.11), with $n = 2{,}000$ neutral simulations and $\alpha' = 0.05$, $\alpha' n = 100$ detections are expected among the neutral simulations. 28 neutral simulations had a $Z$ above the threshold and 96 had exactly the threshold value. Counting only the 28 would give a false-positive rate of 1.4%, and counting all 124 a rate of 6.2%. With the rule, $\gamma = (100 - 28)/96 = 0.75$, and $28 + 0.75 \times 96 = 100$ neutral simulations, exactly 5%, count as detections. The rule changes power in the same way. Among the 1,000 selection simulations of this run, 768 had a $Z$ above the threshold and 3 had exactly the threshold value, so power was 76.8% when only the simulations strictly above the threshold were counted, 77.0% with the rule ($768 + 0.75 \times 3 = 770.25$) and 77.1% when all tied simulations were counted; the threshold, 1.633, was the same in the three cases. Measured on 100,000 independent neutral simulations, this threshold gave 4.85% false positives with the rule, against 1.65% when only simulations strictly above it were counted.

*Effect.* When no neutral simulation has the same Z as the threshold, this rule makes no difference; for example, for the trend test in the reference case, no other neutral simulation had the same Z as the threshold. When ties occur, the false-positive rate is close to 5% (Section S2), and power is higher than when only simulations strictly above the threshold are counted: by 0.2 percentage points in the example above, and by up to 4 points when ties were frequent and selection weak (haploid populations with $p_0 = 0.01$; Section S6).

#### *S1.11 | The sampling-only threshold*

Besides the drift-aware threshold, SPOT reports the results of the sampling-only threshold (Section 2.5 of the main text). This threshold is not set from simulations but from a formula: $z_{1-\alpha'}$, the value exceeded with probability $\alpha'$ by a variable that follows a standard normal distribution (1.645 for $\alpha' = 0.05$). It assumes that, without selection, $Z$ follows a standard normal distribution, which holds approximately when the sampled gene copies are independent draws from a population whose allele frequency does not change, so that the sampling of gene copies is the only source of chance. SPOT applies this threshold to the same neutral and selection simulations as the drift-aware threshold, and reports, for the trend test, the proportion of neutral simulations above it (its actual false-positive rate) and the proportion of selection simulations above it (the power it would report). A simulation counts as a detection only if its $Z$ is strictly above $z_{1-\alpha'}$.

*Why the two thresholds differ.* The neutral simulations include every source of chance of the design: drift, related individuals and errors in genotype calls, as well as the limited number of values that $Z$ can take. The sampling-only threshold ignores all of them. Its false-positive rate is therefore higher than intended when these sources add variation to $Z$ (strong drift, relatives, low depth), and it can be lower than intended when $Z$ takes few values, for example for a rare allele (Section S2).

*Without drift.* With "No drift (infinite population)", the neutral simulations contain no drift, and the drift-aware threshold accounts only for sampling, genotype calling and relatedness.

With unrelated individuals and perfect genotypes, it then rests on the same assumptions as the sampling-only threshold, and the two thresholds are close but not identical: the formula assumes that $Z$ follows a normal distribution, whereas the neutral simulations give the actual distribution of $Z$, including the limited number of values it can take. For example, in a diploid population without drift, with $p_0 = 0.01$, $s = 0.05$, the sampling design of the reference case, unrelated individuals and perfect genotypes (the example without drift), the drift-aware threshold of the trend test was 1.633 instead of 1.645. Measured on 100,000 independent neutral simulations, it gave 4.85% false positives, against 1.65% for the sampling-only threshold, and the powers reported by the two thresholds were 77.0% and 76.4%.

#### *S1.12 | Number of neutral simulations, and many loci*

To place the threshold, about ten neutral simulations should lie above it. This number is a compromise between precision and speed: with fewer simulations above the threshold, its position would vary much more between runs, and with many more, calculations with many loci would become too slow. With this rule, the realised false-positive rate stayed close to the intended rate up to 1,000 loci (Section S2.3; Table S3). SPOT therefore runs the larger of two numbers of neutral simulations: twice the number of selection simulations ("Selection simulations (at least twice as many neutral)"), or $10/\alpha'$, that is, max(2 × number of selection simulations, $10/\alpha'$). For example, with $\alpha = 0.05$ and 100 candidate loci ("Candidate loci tested, L (each tested at α′ = α/L)"), $\alpha' = 0.0005$ and SPOT runs 10/0.0005 = 20,000 neutral simulations. This number is capped at 200,000 for a single calculation, which is reached with 1,000 loci at $\alpha = 0.05$, and at 20,000 per design for the power curve, the sample-size planner and the comparison of sexes.

Beyond these caps, too few neutral simulations lie above the threshold to place it. SPOT then fits a normal distribution to the neutral values of $Z$, sets the threshold at their mean plus $z_{1-\alpha'}$ times their standard deviation (where $z_{1-\alpha'}$ is the $1 - \alpha'$ quantile of the standard normal distribution). This fit is poorly calibrated, because the distribution of $Z$ is not normal in its upper tail. SPOT then shows the note "Unreliable threshold." and explains, under the setting of the number of loci, that even the maximum number of neutral simulations leaves too few of them beyond the threshold, so that the threshold is extrapolated with a normal (Gaussian) fit, which can be far off. Its calibration is described in Section S2.3, and results with more than 1,000 loci should therefore be taken only as an indication.

#### *S1.13 | Other outputs*

*Loss of genetic diversity.* SPOT reports, under "Genetic diversity expected to be lost between the first and the last point" in the interface, the expected proportion of neutral genetic diversity lost by drift between the first and last sampling dates. Genetic diversity is measured by $2p(1 - p)$, where $p$ is the frequency of an allele, and written $H_g$ in generation $g$. In each generation, drift keeps on average a proportion $1 - 1/(2N_e)$ of the diversity of the previous generation in diploids and haplodiploids, which carry $2N_e$ gene copies in an *ideal population of size* $N_e$, and $1 - 1/N_e$ in haploids, which carry $N_e$ gene copies, $N_e$ being then the number of breeding individuals (Wright, 1931; Crow & Kimura, 1970). In diploids and haplodiploids, for example, $H_g = H_{g-1} \times (1 - 1/(2N_{e,g}))$, where $N_{e,g}$ *is the value of* $N_e$ *in generation* $g$, given by its breeding numbers (Section S1.5). Applying this step to each of the $G$ generations between the first and last sampling dates multiplies these proportions, so that the proportion of genetic diversity lost, $1 - H_G/H_0$, is given by Equation (5) in diploids and haplodiploids and by Equation (6) in haploids.

$$1 - \frac{H_G}{H_0} = 1 - \prod_{g=1}^{G}\left(1 - \frac{1}{2N_{e,g}}\right) \tag{5}$$

$$1 - \frac{H_G}{H_0} = 1 - \prod_{g=1}^{G}\left(1 - \frac{1}{N_{e,g}}\right) \tag{6}$$

*With migration.* When migration is set (Section S1.6), migrants bring back diversity every generation: diversity is lost more slowly and finally levels off instead of declining to zero. SPOT then follows, in each generation, drift and then migration, as in its simulations. In diploids and haplodiploids,

$$H_g = (1 - m)^2\left(1 - \frac{1}{2N_{e,g}}\right)H_{g-1} + \left[1 - (1 - m)^2\right]H_0 \tag{7}$$

with $1/N_{e,g}$ in place of $1/(2N_{e,g})$ in haploids, and SPOT reports $1 - H_G/H_0$. Two gene copies taken at random both come from the population with probability $(1 - m)^2$; their diversity is then that of the previous generation after drift (first term of Equation 7). Otherwise, at least one of them comes from a migrant, and their diversity is that of the source, assumed equal to the diversity at the first sampling date, $H_0$ (second term in (7); continent–island model, Wright, 1931). With $m = 0$, Equation (7) gives Equations (5) and (6). This calculation concerns the neutral genome, so it does not depend on the frequency of the favoured allele in the source. For example, in a diploid population with $N_e$ = 100, 90 generations remove 36% of the diversity without migration, 18% with $m$ = 0.01 and 4% with $m$ = 0.05.

*Expected final frequency without drift.* SPOT also reports, in the text under the trajectory chart of the interface, the frequency of the favoured allele at the last sampling date if there were no drift, obtained by running the same model with infinite breeding numbers (no drift step).

*Allele lost or fixed by drift.* When, in more than half of the neutral simulations, drift has lost or fixed the allele in the population by the last date, SPOT reports this proportion under the main power result, and, when power is below 50%, adds that drift has removed the variation the test needs.

### S2 | False-positive rate of the drift-aware threshold

This section details the analysis of Section 2.8.1 of the main text, which measured how often the drift-aware threshold declares selection when there is none.

#### *S2.1 | Procedure*

The false-positive rate of a threshold cannot be measured on the neutral simulations that were used to set it, because these simulations have, by construction, a false-positive rate of exactly 5% (Section S1.10). It must be measured on other neutral simulations, generated independently with the same settings. For each test case, we proceeded as follows.

(1) We ran SPOT 20 times with different random seeds (1,000–1,019), each time with the default 1,000 selection simulations and 2,000 neutral simulations, and recorded the 20 thresholds of each test (trend and first-versus-last). Each threshold is the one a user would obtain from one run of SPOT.

(2) We ran 100,000 further neutral simulations with the same settings and another seed.

(3) For each of the 20 thresholds, the realised false-positive rate is the proportion of these 100,000 neutral simulations whose $Z$ is above the threshold, with simulations whose $Z$ equals the threshold counted as a fraction of a detection, as in SPOT (Section S1.10). It estimates the probability that a neutral locus is declared under selection with this threshold.

(4) We report the mean, the standard deviation, the minimum and the maximum of the 20 realised rates.

(5) We applied the sampling-only threshold (1.645 for $\alpha = 0.05$ and one locus; Section S1.11) to the same 100,000 neutral simulations. For example, in the reference case, 10.3% of these neutral simulations had a $Z$ above 1.645 for the trend test, about twice the intended 5%, whereas the 20 drift-aware thresholds gave 5.00% on average (Table S2).

#### *S2.2 | Test cases*

All test cases use the settings of the reference case unless stated otherwise: a haplodiploid population sampled in 1935, 1955, 1975 and 2025 (generations 0, 20, 40 and 90; generation time 1 year), with 20, 20, 20 and 40 females; mean depth 10× at the earlier dates and 15× at the last; per-read error rate 0.005; minimum depth 3; $p_0$ = 0.1; unrelated individuals; a constant population; no migration; $N_f = N_m$ = 1,000, *that is,* $N_e$ = *1,500.*

In the 24 test cases of Table S1, *the inheritance system and* $N_e$ *were changed*, to vary the amount of drift: each of the three inheritance systems was *combined with eight values of* $N_e$*, 100*, 300, 1,000, 3,000, 10,000, 30,000, 100,000 and 1,000,000. *The* $N_e$ *of the reference case, 1,500, is* not among them, so the reference case appears only in Table S2, and the 36 test cases are all different. The breeding numbers *giving each* $N_e$ *were* $N = N_e$ in haploids, $N_f = N_m = N_e/2$ in diploids and $N_f = N_m = N_e/1.5$ in haplodiploids (Section S1.5).

In the 11 test cases of Table S2, one setting of the reference case was changed at a time: two sampling dates (1935 and 2025, with 20 and 40 females); ten sampling dates every 10 years from 1935 to 2025, with 20 females each; $p_0$ = 0.01 or 0.5; a mean depth of 2× at all dates (minimum depth 1); perfect genotypes; four siblings per family; a ten-fold exponential decline in breeding numbers by the last date; a bottleneck *(*$N_e$ = *1,500, 100 and 1,500 in* the three intervals); migration of 1% per generation from a source with frequency $p_0$; or 200 females at every date.

**Table S1.** Realised false-positive rate (%) for 24 combinations of *inheritance system and* $N_e$*, design* of the reference case otherwise, at an intended rate of 5%.

| Inheritance | $N_e$ | Trend test: drift-aware threshold | Trend test: sampling-only threshold | First-versus-last: drift-aware threshold | First-versus-last: sampling-only threshold |
|---|---|---|---|---|---|
| Haploid | 100 | 5.27 ± 0.41 (4.7–6.0) | 13.58 | 5.14 ± 0.38 (4.4–5.8) | 14.38 |
| Haploid | 300 | 4.95 ± 0.54 (3.7–6.2) | 15.77 | 4.88 ± 0.42 (4.3–6.0) | 14.67 |
| Haploid | 1,000 | 4.83 ± 0.35 (4.1–5.3) | 12.02 | 4.87 ± 0.43 (4.2–5.6) | 10.03 |

| Inheritance | $N_e$ | Trend test: drift-aware threshold | Trend test: sampling-only threshold | First-versus-last: drift-aware threshold | First-versus-last: sampling-only threshold |
|---|---|---|---|---|---|
| Haploid | 3,000 | 5.03 ± 0.41 (4.1–5.6) | 8.12 | 5.06 ± 0.53 (4.2–6.3) | 6.96 |
| Haploid | 10,000 | 4.85 ± 0.50 (3.9–5.6) | 6.02 | 4.95 ± 0.39 (4.6–6.1) | 5.42 |
| Haploid | 30,000 | 4.98 ± 0.46 (4.0–5.6) | 5.37 | 4.89 ± 0.38 (4.4–6.0) | 4.98 |
| Haploid | 100,000 | 5.05 ± 0.47 (4.1–5.7) | 5.15 | 4.96 ± 0.39 (4.5–6.0) | 4.86 |
| Haploid | 1,000,000 | 4.99 ± 0.46 (4.0–6.1) | 5.04 | 4.97 ± 0.37 (4.6–5.9) | 4.81 |
| Diploid | 100 | 4.85 ± 0.44 (3.9–5.6) | 18.36 | 4.92 ± 0.45 (4.3–5.8) | 18.60 |
| Diploid | 300 | 5.08 ± 0.55 (3.9–6.1) | 18.34 | 4.96 ± 0.45 (3.9–5.5) | 16.27 |
| Diploid | 1,000 | 5.01 ± 0.47 (4.2–6.0) | 12.08 | 5.00 ± 0.45 (4.0–5.9) | 9.71 |
| Diploid | 3,000 | 5.17 ± 0.32 (4.3–5.6) | 7.89 | 5.21 ± 0.53 (3.9–6.1) | 6.11 |
| Diploid | 10,000 | 5.13 ± 0.44 (4.3–5.8) | 5.72 | 5.23 ± 0.48 (4.2–5.9) | 4.43 |
| Diploid | 30,000 | 5.20 ± 0.42 (4.2–5.9) | 5.07 | 5.21 ± 0.61 (4.0–6.0) | 3.98 |
| Diploid | 100,000 | 5.15 ± 0.48 (4.2–6.2) | 4.82 | 5.19 ± 0.59 (4.0–6.0) | 3.83 |
| Diploid | 1,000,000 | 5.15 ± 0.50 (4.3–6.5) | 4.72 | 5.17 ± 0.61 (4.0–6.1) | 3.78 |
| Haplodiploid | 100 | 5.02 ± 0.59 (3.8–6.1) | 18.21 | 4.84 ± 0.60 (3.8–6.1) | 18.41 |
| Haplodiploid | 300 | 4.86 ± 0.31 (4.3–5.4) | 18.05 | 4.98 ± 0.41 (4.3–5.6) | 16.21 |
| Haplodiploid | 1,000 | 5.09 ± 0.54 (3.9–6.0) | 12.36 | 5.17 ± 0.51 (4.3–6.0) | 9.94 |
| Haplodiploid | 3,000 | 5.08 ± 0.38 (4.3–5.7) | 7.94 | 5.16 ± 0.54 (3.7–6.0) | 6.11 |
| Haplodiploid | 10,000 | 5.18 ± 0.47 (4.2–5.9) | 5.79 | 5.23 ± 0.47 (4.2–5.8) | 4.46 |
| Haplodiploid | 30,000 | 5.20 ± 0.42 (4.5–6.0) | 5.06 | 5.25 ± 0.60 (4.1–6.0) | 3.98 |
| Haplodiploid | 100,000 | 5.15 ± 0.43 (4.4–6.2) | 4.83 | 5.21 ± 0.60 (4.0–5.9) | 3.82 |
| Haplodiploid | 1,000,000 | 5.14 ± 0.49 (4.3–6.5) | 4.72 | 5.18 ± 0.62 (4.0–6.2) | 3.78 |

Drift-aware threshold: mean ± standard deviation (minimum–maximum) of the realised rates of 20 thresholds, each set from 2,000 neutral simulations. All rates measured on 100,000 independent neutral simulations.

**Table S2.** Realised false-positive rate (%) for the reference case and 11 modified versions of it, each differing in one setting (haplodiploid, $N_e$ = 1,500), at an intended rate of 5%.

| Test case | Trend test: drift-aware threshold | Trend test: sampling-only threshold | First-versus-last: drift-aware threshold | First-versus-last: sampling-only threshold |
|---|---|---|---|---|
| Reference case | 5.00 ± 0.47 (4.0–6.2) | 10.33 | 5.09 ± 0.50 (4.3–5.9) | 8.15 |
| Two dates (1935, 2025) | 5.08 ± 0.48 (4.1–6.2) | 8.20 | 5.08 ± 0.48 (4.1–6.2) | 8.20 |
| Ten dates, 20 individuals each | 5.09 ± 0.55 (4.2–6.3) | 12.03 | 4.99 ± 0.44 (4.1–5.9) | 8.36 |
| $p_0$ = 0.01 | 4.92 ± 0.55 (3.9–5.9) | 4.49 | 5.00 ± 0.50 (4.3–6.0) | 1.69 |
| $p_0$ = 0.5 | 5.17 ± 0.65 (4.0–6.7) | 12.49 | 5.12 ± 0.69 (4.3–7.3) | 11.57 |
| Mean depth 2× at all dates | 5.01 ± 0.58 (4.2–6.2) | 12.78 | 5.05 ± 0.56 (4.2–5.8) | 11.52 |
| Perfect genotypes | 5.07 ± 0.39 (4.4–5.7) | 11.00 | 5.12 ± 0.33 (4.5–5.7) | 8.52 |
| Four siblings per family | 4.98 ± 0.47 (4.1–5.8) | 19.45 | 4.94 ± 0.53 (4.0–5.8) | 19.65 |
| Decline to 10% by the last date | 5.00 ± 0.48 (4.1–5.9) | 18.74 | 5.09 ± 0.40 (4.5–5.9) | 15.10 |
| Bottleneck ($N_e$ 1,500, 100, 1,500) | 4.99 ± 0.52 (3.6–6.0) | 17.42 | 5.00 ± 0.63 (3.5–6.2) | 15.41 |
| Migration, 1% per generation | 5.26 ± 0.44 (4.5–6.0) | 8.06 | 5.25 ± 0.55 (4.1–6.4) | 6.04 |
| 200 individuals per date | 5.13 ± 0.62 (3.9–5.9) | 23.28 | 5.06 ± 0.68 (3.9–6.5) | 22.37 |

#### *S2.3 | Many loci*

When $L$ candidate loci are tested, each locus is tested at $\alpha' = 0.05/L$, and the threshold lies much further in the tail of the neutral distribution of $Z$. For four of the test cases above (Section 2.8.1 of the main text), we repeated the analysis with $L$ = 10, 100 and 1,000 candidate loci, that is, $\alpha'$ = 0.005, 0.0005 and 0.00005. SPOT then ran 2,000, 20,000 and 200,000 neutral simulations per threshold (Section S1.12), and the realised rates were measured on 100,000 ($L$ = 1 and 10) or 1,000,000 ($L$ = 100 and 1,000) independent neutral simulations, so that enough of them lie above the threshold. For comparison, we repeated the analysis with the threshold set from a normal distribution fitted to 2,000 neutral simulations, to assess whether such a fit could replace the additional neutral simulations (Section 2.8.1 of the main text). Table S3 gives, for each threshold, the ratio of the realised to the intended false-positive rate: a ratio of 1 means that the threshold has exactly the intended rate, and a ratio of 2 that it gives twice as many false positives as intended.

*Beyond 1,000 loci.* With more than 1,000 loci at $\alpha$ = 0.05, SPOT sets the threshold from a normal distribution fitted to 200,000 neutral simulations (Section S1.12). To measure its calibration, we applied this threshold with 2,000 and 10,000 loci in the three inheritance systems (design of the reference case, $N_e$ = 1,500, $s$ = 0.05 for power) and measured its false-positive rate on 2,000,000 independent neutral simulations. For a common allele ($p_0$ = 0.1), it gave 0.2–0.7 times the intended rate for the trend test and almost no false positives for the first-versus-last test, so that power was underestimated (for example, 52% instead of 81% for the haplodiploid first-versus-last test with 10,000 loci). For a rare allele ($p_0$ = 0.01), it gave 7.4–118.5 times the intended rate for the trend test, and the power of this test was overestimated (for example, 59% instead of 47% for haploids with 2,000 loci).

**Table S3.** Ratio of the realised to the intended false-positive rate when $L$ candidate loci are tested at $\alpha' = 0.05/L$ (haplodiploid, design of the reference case).

| Test case | $L$ | $\alpha'$ | Neutral simulations per threshold | Independent neutral simulations | Trend: drift-aware threshold | Trend: normal fit to 2,000 | Trend: sampling-only | First-versus-last: drift-aware threshold | First-versus-last: normal fit to 2,000 | First-versus-last: sampling-only |
|---|---|---|---|---|---|---|---|---|---|---|
| $N_e = 300$ | 1 | 0.05 | 2,000 | 100,000 | 0.97 (0.86–1.09) | – | 3.61 | 1.00 (0.86–1.13) | – | 3.24 |
| | 10 | 0.005 | 2,000 | 100,000 | 0.96 (0.65–1.39) | – | 20.69 | 1.00 (0.38–1.90) | – | 14.79 |
| | 100 | 0.0005 | 20,000 | 1,000,000 | 1.11 (0.58–1.67) | 2.55 | 124.77 | 1.05 (0.61–1.78) | 0.74 | 70.40 |
| | 1000 | 0.00005 | 200,000 | 1,000,000 | 1.06 (0.66–1.54) | 2.46 | 762.58 | 1.10 (0.55–1.74) | 0.59 | 346.84 |
| Reference case ($N_e$ = 1,500) | 1 | 0.05 | 2,000 | 100,000 | 1.00 (0.80–1.25) | – | 2.07 | 1.02 (0.85–1.18) | – | 1.63 |
| | 10 | 0.005 | 2,000 | 100,000 | 1.22 (0.63–2.02) | – | 5.52 | 1.18 (0.51–1.75) | – | 2.59 |
| | 100 | 0.0005 | 20,000 | 1,000,000 | 1.19 (0.42–2.48) | 0.95 | 14.49 | 1.08 (0.67–1.51) | 0.07 | 2.97 |
| | 1000 | 0.00005 | 200,000 | 1,000,000 | 1.05 (0.68–1.70) | 0.85 | 39.06 | 1.06 (0.68–1.43) | 0.02 | 3.96 |
| $N_e$ = 100,000 | 1 | 0.05 | 2,000 | 100,000 | 1.03 (0.88–1.23) | – | 0.96 | 1.04 (0.80–1.18) | – | 0.76 |
| | 10 | 0.005 | 2,000 | 100,000 | 1.22 (0.67–1.79) | – | 0.88 | 1.01 (0.57–1.44) | – | 0.37 |
| | 100 | 0.0005 | 20,000 | 1,000,000 | 1.12 (0.59–1.81) | 0.84 | 0.76 | 1.06 (0.47–1.55) | 0.01 | 0.00 |
| | 1000 | 0.00005 | 200,000 | 1,000,000 | 0.98 (0.32–1.48) | 0.47 | 0.40 | 1.02 (0.53–1.64) | 0.00 | 0.00 |
| Reference case, $p_0$ = 0.01 | 1 | 0.05 | 2,000 | 100,000 | 0.98 (0.78–1.18) | – | 0.90 | 1.00 (0.87–1.19) | – | 0.34 |
| | 10 | 0.005 | 2,000 | 100,000 | 1.06 (0.51–1.68) | – | 2.00 | 1.17 (0.44–1.75) | – | 0.15 |
| | 100 | 0.0005 | 20,000 | 1,000,000 | 1.00 (0.70–1.55) | 4.84 | 3.37 | 0.98 (0.70–1.73) | 0.13 | 0.02 |
| | 1000 | 0.00005 | 200,000 | 1,000,000 | 1.05 (0.57–1.53) | 8.31 | 6.28 | 1.02 (0.76–1.58) | 0.08 | 0.02 |

Drift-aware threshold: mean (minimum–maximum) over 20 thresholds. Normal fit: mean over 20 thresholds from a normal distribution fitted to 2,000 neutral simulations; with $L$ = 1 and 10, 2,000 neutral simulations suffice and the two rules coincide (–). A ratio of 1 is the intended rate.

### S3 | Agreement with theoretical expectations

This section checks that SPOT simulates selection, sampling and drift as intended, by comparing its simulations with results that can be calculated without simulation: the change in allele frequency under selection without drift or with weak drift (Section S3.1), the exact power without drift (Section S3.2; Figure S1a) and the loss of genetic diversity under drift (Section S3.3; Figure S1b). We also checked that sampling unrelated individuals through the procedure used for families (families of one individual) gives the same power as the default procedure (Section S3.4).

#### *S3.1 | Selection*

*Selection without drift.* Without drift ("No drift (infinite population)"), the frequency of the favoured allele changes deterministically, as given by Equations (1) and (2) (Section S1.3) applied in every generation. For haploids, these generations can be combined into a closed form: after *t* generations of selection, the frequency is

$$p_t = \frac{p_0(1+s)^t}{1-p_0+p_0(1+s)^t} \tag{8}$$

We implemented these equations and compared SPOT's trajectories with them over 300 generations, for all combinations of 6 inheritance systems and dominance coefficients (haploid; diploid with $h$ = 0, 0.5 and 1; haplodiploid with $h$ = 0 and 0.5, with the same selection in both sexes), 3 selection coefficients ($s$ = 0.01, 0.05 and 0.2) and 2 starting frequencies ($p_0$ = 0.01 and 0.2), that is, 6 × 3 × 2 = 36 trajectories. The largest difference was $2 \times 10^{-15}$, which is the rounding error of the computer.

*Selection with weak drift.* This check tests whether SPOT simulates selection correctly when drift acts as well. Drift is random and should not push the allele frequency in any systematic direction, so the mean of many simulated trajectories should follow the trajectory without drift; a systematic difference would reveal an error. We simulated 500 trajectories of 300 generations in a diploid population with $N_f = N_m = 10^6$ breeding adults ($N_e = 2 \times 10^6$), $h$ = 0.5, $s$ = 0.05 and $p_0$ = 0.01. In this population, more than 20 copies of the rarer allele are always expected, so SPOT draws drift from a normal distribution (Section S1.4). The population had to be very large: with strong drift, the mean of the trajectories differs from the trajectory without drift even when the simulation is correct, because selection changes the frequency non-linearly, so this check is only meaningful when drift is weak. The favoured allele rose from 0.01 to 0.94 over the 300 generations, and at every generation the mean of the 500 trajectories was within 0.0006 of the trajectory without drift. The largest difference (0.4959 against 0.4953) occurred at generation 186, when the frequency was about 0.5: there the trajectory rises fastest and drift is strongest, so small random differences between trajectories are expected to be the most visible.

#### *S3.2 | Power without drift*

Without drift and with perfect genotypes, the only source of chance is the sampling of individuals. Power can then be calculated exactly: the numbers of copies of the favoured allele in the samples of the different dates are independent binomial variables, whose probabilities follow from the genotype frequencies after selection in each sampled generation (Equations 1 and 2), so every possible combination of counts can be listed with its probability. There are, for instance, up to 5.6 million combinations for four sampling dates. The exact power is the total probability of the combinations whose $Z$ exceeds the threshold, with combinations whose $Z$ equals the threshold counted with the weight $\gamma$ of Section S1.10.

We made this comparison for all combinations of 5 inheritance systems and dominance coefficients (haploid; diploid with $h$ = 0, 0.5 and 1; haplodiploid with $h$ = 0.5, females sampled), 2 starting frequencies ($p_0$ = 0.05 and 0.2), 4 selection coefficients ($s$ = 0.005, 0.01, 0.02 and 0.05) and 2 sampling designs (four dates, 1935, 1955, 1975 and 2025, with 20, 20, 20 and 40 individuals; or two dates, 1935 and 2025, with 20 and 40), that is, 5 × 2 × 4 × 2 = 80 designs. SPOT used 5,000 selection and 10,000 neutral simulations per design. For each design and each test (trend and first-versus-last), we compared SPOT's power with the exact power for two thresholds: the sampling-only threshold, and SPOT's drift-aware threshold, for which the exact power was calculated at the same threshold and with the same weight for ties (240 comparisons in total, 120 for each threshold).

For the drift-aware threshold, SPOT's power and the exact power differed by 0.00 percentage points on average and by at most 1.4 points; 2 of the 120 differences exceeded two standard errors (against about 6 expected by chance; Figure S1a). The exact false-positive rate of SPOT's thresholds, calculated from the exact distribution of $Z$, was 5.01% on average (4.43–5.73%) for an intended 5%: the remaining spread comes from placing each threshold with a finite number of neutral simulations (Section S1.12). The power of the sampling-only threshold also agreed with the exact calculation within the variation expected between runs.

#### *S3.3 | Drift: loss of genetic diversity*

Without selection, drift reduces the genetic diversity of a population, measured by $2p(1-p)$, where $p$ is the frequency of an allele. In expectation, the proportion of diversity lost after $G$ generations is given by Equations (5) and (6) (Section S1.13), where $N_{e,g}$ *is the value of* $N_e$ *in generation* $g$: $4N_fN_m/(N_f + N_m)$ in diploids, $9N_fN_m/(4N_m + 2N_f)$ in haplodiploids, and $N$ in haploids (Section S1.5).

We ran 100,000 neutral simulations of 90 generations for each of nine scenarios (Table S4), with $p_0 = 0.3$, and calculated, in every generation, the proportion of diversity lost, that is, one minus the mean of $p(1-p)$ over the simulations divided by its value at generation 0. This proportion was compared with the theoretical value in every generation (Figure S1b).

In all scenarios and at every generation, the simulated proportion lost differed from the theoretical value by at most 0.0042. At generation 90, the differences ranged from −1.7 to +1.4 standard errors (Table S4). The largest difference at any single generation (3.5 standard errors) occurred in the diploid scenario with very unequal numbers of breeding females and males ($N_f = 100$, $N_m = 20$). The loss of genetic diversity reported by SPOT (Section S1.13) is not estimated from simulations: it is calculated with Equations (5) and (6). For the nine scenarios, the loss reported by SPOT differed from the theoretical value of Table S4 by less than 0.0001, which confirms that this output is calculated correctly.

*With migration.* We repeated this comparison against Equation (7) in eight scenarios with migration: diploid, haploid and haplodiploid populations, one with $N_e$ set per interval, $m$ from 0.001 to 0.05, and migrants carrying the starting frequency $p_0$. At every generation, the simulated proportion lost differed from Equation (7) by at most 0.0024, and at generation 90 by −1.8 to +1.7 standard errors. The loss reported by SPOT is calculated with Equation (7).

**Table S4.** Proportion of genetic diversity lost after 90 generations of drift ($p_0 = 0.3$; 100,000 neutral simulations per scenario).

| Scenario | Simulated (± standard error) | Theoretical |
|---|---|---|
| Haploid, $N = 100$ | 0.5974 ± 0.0015 | 0.5953 |
| Haploid, $N = 1{,}000$ | 0.0862 ± 0.0008 | 0.0861 |
| Diploid, $N_f = N_m = 50$ | 0.3639 ± 0.0014 | 0.3631 |
| Diploid, $N_f = N_m = 500$ | 0.0442 ± 0.0006 | 0.0440 |
| Diploid, $N_f = 100$, $N_m = 20$ | 0.4895 ± 0.0015 | 0.4921 |

| Scenario | Simulated (± standard error) | Theoretical |
|---|---|---|
| Haplodiploid, $N_f = N_m = 100$ | 0.2603 ± 0.0013 | 0.2596 |
| Haplodiploid, $N_f = 300$, $N_m = 50$ | 0.2332 ± 0.0012 | 0.2344 |
| Diploid, $N_f = N_m = 500$, decline to 10% | 0.1601 ± 0.0011 | 0.1597 |
| Diploid, $N_e$ = 1,000, 50 and 1,000 in the three intervals | 0.2114 ± 0.0012 | 0.2102 |

#### *S3.4 | Sampling of relatives*

SPOT generates sampled individuals by two procedures (Section S1.7). In the procedure for unrelated individuals, the default, the genotype of each sampled individual is drawn directly from the genotype frequencies of the offspring after selection. In the procedure for families, a mother and a father are drawn from the genotype frequencies of the female and male offspring after selection, and each sibling then receives one allele from each parent, following Mendel's rules, without further selection. The genotypes of siblings therefore follow the allele frequencies of their selected parents, rather than exactly the genotype frequencies of the offspring after selection. To check that this difference does not by itself change power, we generated unrelated individuals with the procedure for families, as families of one individual, and compared the power obtained with that of the procedure for unrelated individuals. Because both procedures then give unrelated individuals, a difference in power could come only from the procedure itself, not from relatedness. We used all combinations of 2 inheritance systems (diploid and haplodiploid), *3 values of* $N_e$ *(300,* 1,500 and 100,000) and 2 selection coefficients ($s$ = 0.02 and 0.05), each run with 3 random seeds, that is, 2 × 3 × 2 × 3 = 36 runs, with the design of the reference case otherwise ($p_0$ = 0.1; 20, 20, 20 and 40 females sampled in 1935, 1955, 1975 and 2025) and perfect genotypes. Each run used 20,000 selection and 40,000 neutral simulations with each procedure, and each procedure set its own threshold from its neutral simulations, as SPOT does. To judge whether a difference between the two procedures was larger than expected by chance, we divided it by its standard error, calculated as for the difference between two proportions estimated from 20,000 selection simulations each; a difference larger than two standard errors is then expected in about 5% of the runs. This standard error slightly underestimates the true one, because it ignores the variation of the threshold between runs (Section S5).

Power ranged from about 20% to 100% over the 36 runs. The two procedures gave powers that differed by −1.4 to +1.0 percentage points for the trend test and by −1.6 to +1.8 points for the first-versus-last test (procedure for families minus procedure for unrelated individuals), with mean differences of +0.03 and +0.02 points, and the differences were positive in some runs and negative in others. In the runs in which power was below 100%, 2 of the 31 differences of the trend test and 4 of the 33 differences of the first-versus-last test exceeded two standard errors, against about 1.5 and 1.7 expected by chance. With one individual per family, the family procedure gave the same power as the procedure for unrelated individuals. The way SPOT simulates families therefore does not change power by itself: when several relatives are sampled, any change in power comes from their relatedness.

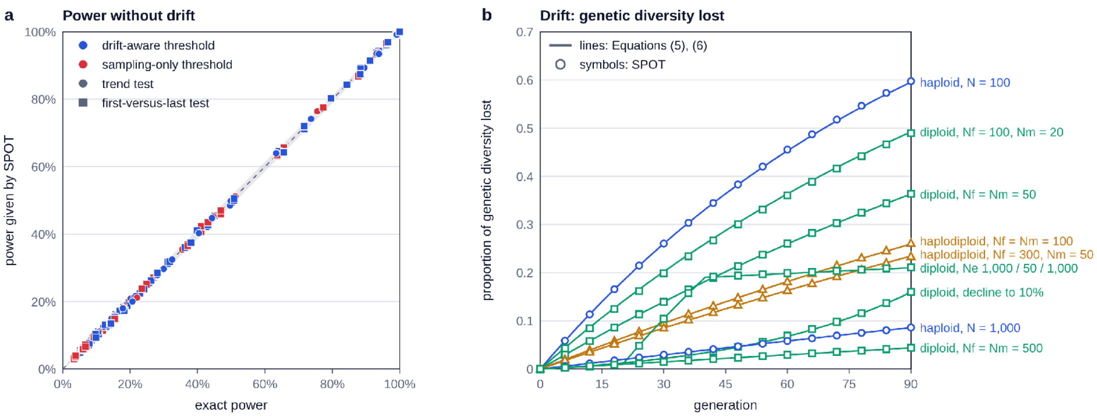


**Figure S1.** Agreement with theoretical expectations (Section S3). (a) Power without drift: power given by SPOT against the exact power, for the 80 designs of Section S3.2, both tests and both thresholds (blue, drift-aware threshold; red, sampling-only threshold; circles, trend test; squares, first-versus-last test, including the designs with two dates); dashed line, 1:1; grey band, ±2 standard errors of a power estimated from 5,000 selection simulations, so that a point inside the band differs from the exact power by no more than expected by chance. (b) Drift: proportion of genetic diversity lost over 90 generations in the nine scenarios of Table S4 ($p_0 = 0.3$); lines, Equations (5) and (6); symbols, mean over 100,000 neutral simulations of SPOT (circles, haploids; squares, diploids; triangles, haplodiploids), whose standard errors (at most 0.0015) are smaller than the symbols. Selection without drift (Section S3.1) agreed with Equation (8) to within the rounding error of the computer and is not shown.

### S4 | Comparison with SLiM

SPOT does not simulate individual organisms. To be fast, it follows the frequency of the favoured allele among breeding females and males (Section S1.3), draws the gene copies of the next breeding adults as binomial samples (Section S1.4), and creates each sampled individual only when it is sampled, from the genotype frequencies of the population (Section S1.7). The aim of this section is to test whether these simplifications change the results, by comparing SPOT with the individual-based simulator SLiM 5.2 (Haller et al., 2026), in which every individual is created, reproduces and can be sampled.

We compared four results of the two programs, for the same scenarios and the same sampling design. (1) Allele frequencies: if SPOT simulates drift and selection as SLiM does, the frequency of the favoured allele should have the same distribution in the two programs at every sampling date. (2) Power, the result that SPOT reports to users, each program setting its own threshold from its own neutral simulations. (3) False-positive rates: if the drift simulated by SPOT is equivalent to that of a population of individuals, a threshold set from SPOT's neutral simulations should give the intended 5% of false positives on SLiM's neutral simulations. (4) Power with SPOT's threshold: the threshold set by SPOT, applied to SLiM's simulations with selection, should give the same power as on SPOT's own simulations. The last two comparisons matter most for users, because they test SPOT as it is used: its threshold and its power, computed from simplified simulations, are meant to apply to populations made of individuals.

#### *S4.1 | SLiM model*

We used SLiM to simulate one locus with two alleles in a Wright–Fisher population of constant size, without mutation or migration. Every individual is simulated. In each generation, SLiM creates each offspring by choosing its mother among the breeding females and its father among the breeding males (in haploids, its single parent among all breeding individuals), each parent with a probability proportional to its fitness; the offspring then inherits one gene copy from each parent, following Mendel's rules. This choice of parents according to their fitness is how SLiM simulates selection in a Wright–Fisher population. SPOT applies selection instead to the survival of offspring before they become breeding adults (Section S1.3). In both programs, a genotype with fitness $1 + s$ therefore contributes on average $1 + s$ times as many gene copies to the next generation as a genotype with fitness 1.

In diploids, $N_f$ females and $N_m$ males had fitness 1, $1 + hs$ and $1 + s$ with zero, one and two copies of the favoured allele. In haploids, $N$ individuals reproduced clonally, with fitness 1 or $1 + s$. SLiM has no haplodiploid inheritance. We therefore gave each haplodiploid male two identical copies of the genome inherited from his mother, by replacing in every generation the copy inherited from his father with a copy of his mother's, and then gave him fitness 1 or $1 + s$ according to this single genome. A father then always transmits his single genome to his daughters, and a son inherits from his mother only, which reproduces haplodiploid inheritance exactly for a single locus.

The favoured allele started at frequency $p_0$ in both sexes (it was given to a proportion $p_0$ of the gene copies of each sex), and selection acted from generation 1, as in SPOT when selection starts at the first sampling date (Section S1.1). At each sampling date, SLiM recorded, in each simulated population, the two quantities that SPOT simulates: the frequency of the favoured allele among breeding females after selection, that is, with each female weighted by her fitness (in haploids, among all breeding individuals); and the number of copies of the favoured allele carried by the sampled females (in haploids, the sampled individuals), each drawn with a probability proportional to her fitness, as SPOT draws sampled individuals from the genotype frequencies after selection. Genotypes were perfect.

#### *S4.2 | Scenarios and comparisons*

*Scenarios.* SPOT and SLiM simulated the same 11 scenarios: six with selection and five without selection (Table S5). The scenarios with selection cover the three inheritance *systems,* $N_e$ *from 200 to 1,500*, starting frequencies of 0.01, 0.05 and 0.1, selection coefficients of 0.02 and 0.05, and additive ($h$ = 0.5) and recessive ($h$ = 0) selection. Each scenario without selection has the settings of the corresponding scenario with selection, with $s$ = 0; the two diploid scenarios with $N_e$ = 1,000 share one. All scenarios used the sampling design of the reference case (samples at generations 0, 20, 40 and 90, with 20, 20, 20 and 40 females; Section S2.2) with perfect genotypes. This comparison therefore concerns drift, selection and sampling, not sequencing and genotype calls, which SPOT simulates after the individuals are sampled (Section S1.8).

*Simulations.* Each scenario, with or without selection, was simulated in 20 runs by each program, with different random seeds, each run giving 2,000 independent simulations (in SLiM, 2,000 populations simulated side by side as separate subpopulations of one run). For the false-positive rates, we used 10,000 SLiM simulations of each scenario without selection (its first run and four further runs of 2,000) and 30,000 further SPOT simulations (20,000 to set a threshold and 10,000 to measure its false-positive rate). The random seeds are given in Section S8.

*Comparisons.* (1) *Allele frequencies.* For each of the 11 scenarios, we compared the 2,000 simulations of one run of each program at generations 20, 40 and 90 (at generation 0, the frequency is $p_0$ in both programs). For the frequency of the favoured allele among breeding females after selection, we compared three statistics: its mean, with the difference between the programs expressed in standard errors; its standard deviation between simulations, as the ratio SPOT/SLiM; and, at generation 90, its whole distribution, with a two-sample Kolmogorov–Smirnov test.

(2) *Power.* In each of the 20 runs of a scenario with selection, each program set the threshold of the trend test from its own 2,000 neutral simulations, with the rule of Section S1.10, and power was the proportion of its own 2,000 selection simulations above this threshold. We compared the mean and the standard deviation of the 20 powers of each program, and expressed the difference between the mean powers in standard errors, calculated from the standard deviations between runs.

(3) *False-positive rates.* For each scenario without selection, a threshold set from 20,000 SPOT neutral simulations was applied to the 10,000 SLiM neutral simulations and to 10,000 independent SPOT neutral simulations. We also applied the sampling-only threshold (Section S1.11) to both sets.

(4) *Power with SPOT's threshold.* For each scenario with selection, a threshold set from 20,000 further SPOT neutral simulations was applied to the selection simulations of both programs in each of the 20 runs, and the mean powers over the 20 runs were compared.

(5) *Regression.* For the mean allele frequencies (1), the mean powers (2) and the false-positive rates of both thresholds (3), we regressed the values given by SLiM on those given by SPOT over all scenarios: a slope of 1 and an intercept of 0 mean that the two programs agree over the whole range of values.

#### *S4.3 | Results*

*Allele frequencies.* At generations 20, 40 and 90, the mean frequency of the favoured allele among breeding females differed between the programs by −1.4 to +2.2 standard errors over the 33 comparisons (11 scenarios × 3 generations), and the ratio of the standard deviations (SPOT/SLiM) was 0.93–1.08. At generation 90, the Kolmogorov–Smirnov test found no difference between the distributions in any scenario ($D = 0.015$–$0.035$, $p = 0.16$–$0.98$).

*Power.* With each program's own threshold, the mean power over 20 runs differed by −0.9 to +1.0 percentage points (SPOT minus SLiM), never by more than two standard errors, and the standard deviation of power between runs was similar in the two programs, 1.3–2.4 points (Table S5; Figure S2a).

*False-positive rates.* The threshold set from 20,000 SPOT neutral simulations gave 4.9–5.5% false positives on SLiM's neutral simulations and 4.9–5.7% on independent SPOT neutral simulations, for an intended 5%. The sampling-only threshold gave 5.4–19.7% on the simulations of both programs (Table S6).

*Power with SPOT's threshold.* Applied to the selection simulations of both programs, SPOT's threshold gave mean powers that differed by −0.4 to +0.3 percentage points between the programs (Table S5).

*Regression.* The slope of the regression of SLiM's values on SPOT's was 0.992 (95% confidence interval 0.980–1.004; $R^2$ = 0.999) for the mean allele frequencies (33 values), 1.017 (0.957–1.078; $R^2$ = 0.998) for the mean powers (6 values) and 1.011 (0.976–1.047; $R^2$ = 0.998) for the false-positive rates of both thresholds (10 values); the 95% confidence intervals of all three intercepts included 0 (Figure S2b–d).

SPOT and SLiM therefore gave the same results in all four comparisons, within the variation expected between simulations: the simplifications of SPOT do not change the allele frequencies, the power or the false-positive rates.

**Table S5.** Scenarios with selection simulated by SPOT and SLiM, and power of the trend test (mean ± standard deviation over 20 runs of 2,000 selection and 2,000 neutral simulations per program).

| Inheritance | $N_f$, $N_m$ ($N_e$) | $p_0$ | $s$ | $h$ | SPOT | SLiM | Difference (SE) | SPOT threshold: on SPOT / on SLiM |
|---|---|---|---|---|---|---|---|---|
| Diploid | 500, 500 (1,000) | 0.1 | 0.02 | 0.5 | 31.8 ± 1.8 | 32.1 ± 1.9 | −0.3 (0.6) | 32.9 / 32.9 |
| Diploid | 500, 500 (1,000) | 0.1 | 0.05 | 0 | 25.2 ± 1.3 | 24.8 ± 1.5 | +0.4 (0.5) | 24.2 / 24.1 |
| Diploid | 100, 100 (200) | 0.1 | 0.05 | 0.5 | 53.4 ± 1.7 | 52.9 ± 2.0 | +0.5 (0.6) | 52.4 / 52.1 |
| Haploid | 1,000 (1,000) | 0.05 | 0.02 | – | 43.8 ± 1.9 | 42.8 ± 1.9 | +1.0 (0.6) | 42.2 / 42.0 |
| Haplodiploid | 500, 500 (750) | 0.1 | 0.02 | 0.5 | 43.1 ± 1.7 | 43.5 ± 2.4 | −0.4 (0.7) | 43.4 / 43.6 |
| Haplodiploid | 1,000, 1,000 (1,500) | 0.01 | 0.05 | 0.5 | 66.3 ± 1.6 | 67.2 ± 1.4 | −0.9 (0.5) | 67.1 / 67.5 |

Power in %. Difference: SPOT minus SLiM, with its standard error. Last column: mean power when a threshold set from 20,000 SPOT neutral simulations of the scenario without selection is applied to the selection simulations of each program. Sampling design of the reference case (20, 20, 20 and 40 females at generations 0, 20, 40 and 90), perfect genotypes.

**Table S6.** False-positive rate of the trend test on the neutral simulations of SPOT and SLiM (%).

| Neutral scenario | SPOT threshold (from 20,000 SPOT simulations) | On 10,000 SLiM simulations | On 10,000 SPOT simulations | Sampling-only threshold, on SLiM | Sampling-only threshold, on SPOT |
|---|---|---|---|---|---|
| Diploid, $N_e = 1{,}000$, $p_0 = 0.1$ | 2.477 | 5.4 | 5.0 | 13.5 | 13.0 |
| Diploid, $N_e = 200$, $p_0 = 0.1$ | 4.240 | 4.9 | 4.9 | 19.7 | 19.7 |
| Haploid, $N = 1{,}000$, $p_0 = 0.05$ | 2.187 | 5.3 | 5.5 | 10.2 | 10.4 |
| Haplodiploid, $N_e = 750$, $p_0 = 0.1$ | 2.654 | 5.5 | 5.7 | 15.4 | 15.2 |
| Haplodiploid, $N_e = 1{,}500$, $p_0 = 0.01$ | 1.735 | 5.0 | 5.0 | 5.4 | 5.4 |

Standard error of each rate on 10,000 simulations: about 0.2 percentage points.

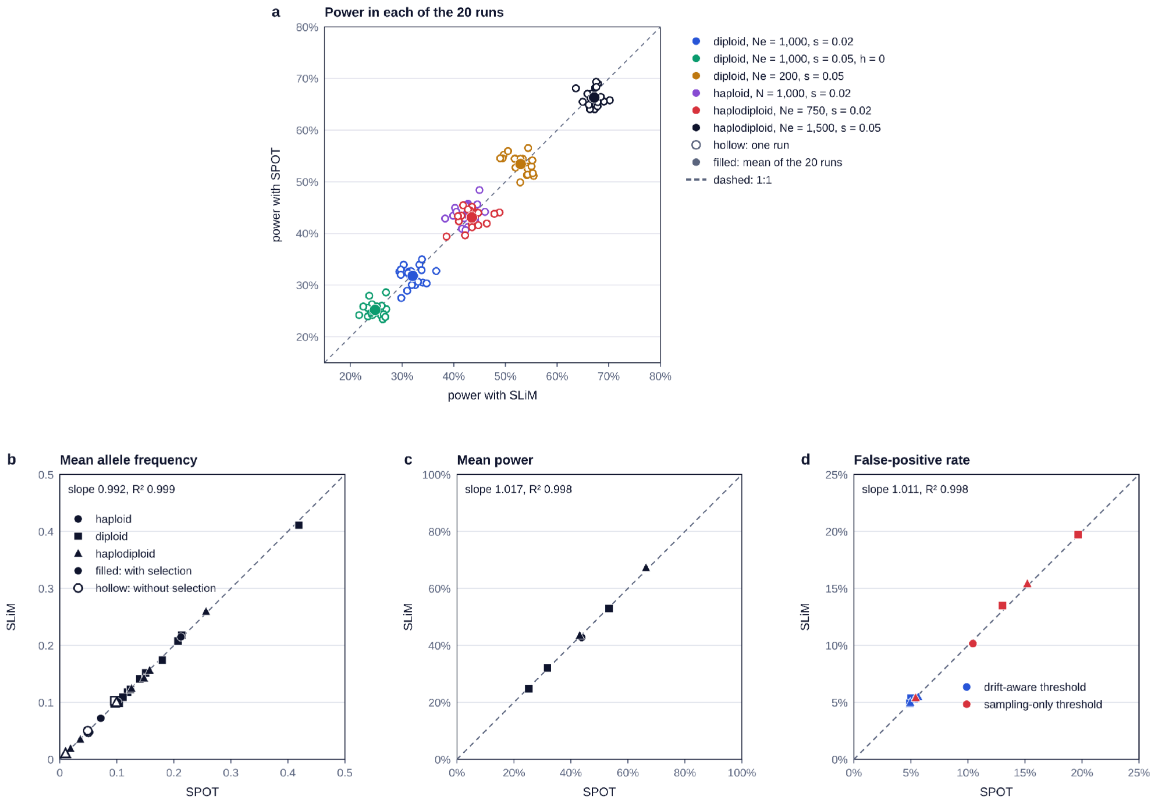


**Figure S2.** Comparison of SPOT with SLiM (Section S4). (a) Power of the trend test in each of the 20 runs of each program (hollow) and mean of the 20 runs (filled), for the six scenarios with selection of Table S5; in each run, each program set its own threshold from its own 2,000 neutral simulations. (b–d) Values given by SLiM against values given by SPOT, with the slope and $R^2$ of the regression of SLiM's values on SPOT's: (b) mean frequency of the favoured allele among breeding females at generations 20, 40 and 90 in the 11 scenarios (filled, with selection; hollow, without selection; shapes, inheritance systems); (c) mean power over the 20 runs, in the six scenarios with selection; (d) false-positive rate of the threshold set from 20,000 SPOT neutral simulations (blue) and of the sampling-only threshold (red), in the five scenarios without selection (Table S6). Dashed lines, 1:1.

### S5 | Precision and speed

SPOT estimates power from a finite number of simulations, so two runs with the same settings and different random seeds never give exactly the same power. This section measures how large this variation between runs is, where it comes from and how it decreases with the number of simulations, and how long a calculation takes. These results matter to users: they tell whether a difference of a few percentage points between two designs is real or comes from the variation between runs, and how many simulations, and how much time, a more precise answer requires.

***S5.1 | Procedure***

*Scenarios.* We used the five example scenarios of the application, "Bumblebees", "Long-lived mammal", "Human ancient DNA", "Bacterial pathogen" and "Large diploid population", whose settings are given in Table S7a. They cover the three inheritance *systems, $N_e$ from 200 to 2* × $10^6$, and series of 7 to 17,000 generations, the longest simulated with rescaling (Section S6).

*Runs.* For each scenario, we ran SPOT 50 times with different random seeds, with 500, 1,000 (the default), 2,000 and 5,000 selection simulations. As in the application, each run used twice as many neutral simulations as selection simulations (Section S1.12), set its own drift-aware threshold from them, and computed the power of the trend test and of the first-versus-last test, with $\alpha = 0.05$ and one locus.

*Variation between runs.* For each scenario, number of selection simulations *R* and test, we calculated the standard deviation of power between the 50 runs.

*Comparison with the binomial standard deviation.* The power of a run varies between runs for two reasons. First, power is the proportion of the R selection simulations whose Z lies above the threshold, and this proportion varies by chance because R is finite. This first source of variation alone has a known size, the binomial standard deviation √(P(1 − P)/R), where P is the power that an infinite number of simulations would give (estimated here by the mean power of the 50 runs). For example, with a power of 50% and R = 1,000, it is √(0.5 × 0.5/1,000) = 0.016, that is, 1.6 percentage points. Second, the threshold is itself set from a finite number of neutral simulations, so it moves slightly from run to run, and each move shifts some selection simulations above or below it. To measure how much this second source adds, we divided the observed standard deviation of power by the binomial standard deviation. A ratio close to 1 means that almost all the variation comes from the finite number of selection simulations; a ratio of 2 means that the movement of the threshold doubles it. This matters for users: when the ratio is large, increasing the number of selection simulations alone does not make power much more precise. The threshold moves most when few neutral simulations have values of Z close to it, and its movement matters most when many selection simulations lie close to it.

*Speed.* We measured the time taken by one complete calculation (simulations, thresholds and power of both tests) with the default numbers of simulations (1,000 selection and 2,000 neutral) and one locus, for the five example scenarios and for the reference case (Section S2.2), and for the reference case with 1,000 candidate loci, which requires 200,000 neutral simulations (Section S1.12). The calculations used the simulation code of SPOT in Node.js (version 22), on one core of an Intel Xeon processor at 2.1 GHz; each calculation was timed five times after a first, untimed run, and we report the median.

#### *S5.2 | Results*

*Variation between runs.* With the default 1,000 selection simulations, the standard deviation of power between runs was 1.5–2.2 percentage points for the trend test and 1.4–3.1 points for the first-versus-last test, depending on the scenario (Table S7b). For example, in "Bumblebees", the mean power of the trend test was 66.6%, its standard deviation 1.5 points, and the 50 runs ranged from 63.2% to 69.7%. The standard deviation decreased as the number of selection simulations increased, approximately in proportion to $1/\sqrt{R}$: with 5,000 selection simulations, it was 0.6–1.1 points for the trend test and 0.6–1.5 points for the first-versus-last test (Figure S3).

*Comparison with the binomial standard deviation.* The observed standard deviation was 0.9–1.8 times the binomial standard deviation for the trend test, and 0.9–2.3 times for the first-versus-last test (Table S7b). For the trend test in "Bumblebees" and "Bacterial pathogen", the ratio was 0.92–1.16: almost all the variation came from the selection simulations. The ratio was largest in "Long-lived mammal", for both tests (1.42–1.80), and in "Large diploid population" for the first-versus-last test (1.76–2.29). For example, in "Large diploid population" with 1,000 selection simulations, the first-versus-last test had a mean power of 66% over the 50 runs; the observed standard deviation was 3.1 points, against 1.5 points for the binomial standard deviation, so the movement of the threshold doubled the variation between runs (single runs ranged from 61% to 73%), not power itself.

*Speed.* With one locus and the default numbers of simulations, a complete calculation took 0.05–0.57 s for the five example scenarios and 0.1 s for the reference case. With 1,000 candidate loci, the reference case took about 7 s.

#### *S5.3 | Practical consequences*

With the default 1,000 selection simulations, the power of the trend test given by a single run is typically within about 3–4.5 percentage points of the value that an infinite number of simulations would give. This precision is enough to tell whether a design has a power of about 30% or about 80%, but not to rank two designs whose powers differ by only a few points, or to decide whether a design whose power is close to a target, for example 80%, reaches it. In these cases, we recommend running SPOT with 5,000 selection simulations, which reduces the standard deviation to 0.6–1.1 points for the trend test and takes roughly four times as long.

Increasing the number of selection simulations alone would not be enough when the threshold varies between runs, as in "Long-lived mammal" and for the first-versus-last test in "Large diploid population": the number of neutral simulations must increase too, so that the threshold is placed more precisely. SPOT does this automatically, because it always runs at least twice as many neutral as selection simulations (Section S1.12). For this reason, the standard deviation decreased with the number of selection simulations even where the threshold added most variation, for example from 3.5 to 1.0 points in "Long-lived mammal" between 500 and 5,000 selection simulations.

**Table S7a.** Settings of the five example scenarios of the application used in Section S5.

| Setting | "Bumblebees" | "Long-lived mammal" | "Human ancient DNA" | "Bacterial pathogen" | "Large diploid population" |
|---|---|---|---|---|---|
| Inheritance | Haplodiploid | Diploid | Diploid | Haploid | Diploid |
| Years per generation | 1 | 20 | 28 | 0.01 | 1 |
| Sampling years | 1935, 1955, 1975, 2025 | 1880, 1950, 2020 | 7000, 5000, 3000 and 1000 BCE; 1000 and 2000 CE | 1850, 1900, 1950, 2000, 2020 | 1900, 1950, 2000, 2020 |
| Generations from the first to the last date | 90 | 7 | 321 | 17,000 (simulated as 1,889 with $\lambda$ = 9) | 120 |
| Individuals per date | 20, 20, 20, 40 | 20, 20, 40 | 50 at each ancient date, 100 present-day | 10, 10, 15, 20, 40 | 30, 30, 30, 60 |
| Breeding numbers ($N_e$) | $N_f = N_m$ = 1,000 (1,500) | $N_e$ = 5,000, then 200 (set per interval) | $N_f = N_m$ = 5,000 (10,000) | $N$ = 100,000 | $N_f = N_m = 10^6$ ($2 \times 10^6$) |
| Starting frequency $p_0$ | 0.01 | 0.1 | 0.1 | 0.05 | 0.05 |
| Selection coefficient $s$ | 0.05 | 0.2 | 0.005 | 0.0002 | 0.02 |
| Mean depth, earlier dates / last date | 10× / 15× | 4× / 15× | 2× / 20× | 10× / 30× | 10× / 15× |
| Per-read error rate (all dates) | 0.005 | 0.005 | 0.01 | 0.005 | 0.005 |
| Minimum depth | 3 | 3 | 1 | 3 | 3 |

All five scenarios: $h$ = 0.5 (no effect in haploids), selection from the first sampling date, unrelated individuals, only females sampled (diploids and haploidiploids), no migration, $\alpha$ = 0.05 and one locus. In "Long-lived mammal", $N_e$ *is set separately* for each interval (Section S1.5): 5,000 between 1880 and 1950 and 200 between 1950 and 2020; 1950 falls 3.5 generations after 1880 and is rounded to generation 4 (Section S1.1). In "Bacterial pathogen", the 17,000 generations are simulated with rescaling (Section S6.1).

**Table S7b.** Variation of power between 50 runs of SPOT with different random seeds, for the five example scenarios of Table S7a (power in %; standard deviations in percentage points).

| Scenario | Selection simulations *R* | Mean power (trend) | SD (trend) | SD expected from *R* (trend) | Ratio (trend) | Range of the 50 runs (trend) | SD (first-versus-last) | Ratio (first-versus-last) |
|---|---|---|---|---|---|---|---|---|
| Bumblebees | 500 | 66.5 | 2.1 | 2.1 | 1.00 | 59.4–71.0 | 2.1 | 0.99 |
| | 1,000 | 66.6 | 1.5 | 1.5 | 1.02 | 63.2–69.7 | 1.4 | 0.96 |
| | 2,000 | 66.6 | 1.1 | 1.1 | 1.06 | 64.0–68.5 | 1.3 | 1.24 |
| | 5,000 | 66.5 | 0.7 | 0.7 | 1.09 | 64.6–67.8 | 1.0 | 1.52 |
| Long-lived mammal | 500 | 24.9 | 3.5 | 1.9 | 1.80 | 18.4–33.4 | 3.0 | 1.58 |
| | 1,000 | 24.7 | 2.2 | 1.4 | 1.61 | 20.3–30.5 | 2.2 | 1.61 |
| | 2,000 | 24.5 | 1.4 | 1.0 | 1.47 | 21.5–27.5 | 1.4 | 1.42 |
| | 5,000 | 24.4 | 1.0 | 0.6 | 1.62 | 21.0–26.6 | 1.0 | 1.68 |
| Human ancient DNA | 500 | 47.4 | 2.6 | 2.2 | 1.17 | 42.6–53.8 | 3.0 | 1.37 |
| | 1,000 | 47.5 | 2.0 | 1.6 | 1.27 | 43.1–51.2 | 1.9 | 1.21 |
| | 2,000 | 47.1 | 1.4 | 1.1 | 1.29 | 43.5–50.1 | 1.4 | 1.31 |
| | 5,000 | 47.1 | 1.1 | 0.7 | 1.56 | 44.9–50.0 | 1.0 | 1.44 |
| Bacterial pathogen | 500 | 72.6 | 2.3 | 2.0 | 1.16 | 68.4–78.6 | 2.1 | 1.05 |
| | 1,000 | 72.6 | 1.6 | 1.4 | 1.10 | 69.7–76.3 | 1.5 | 1.06 |
| | 2,000 | 72.4 | 1.0 | 1.0 | 0.97 | 70.2–74.3 | 1.1 | 1.05 |
| | 5,000 | 72.5 | 0.6 | 0.6 | 0.92 | 71.3–73.9 | 0.6 | 0.92 |
| Large diploid population | 500 | 73.2 | 2.6 | 2.0 | 1.33 | 66.8–79.0 | 3.7 | 1.76 |
| | 1,000 | 73.3 | 1.7 | 1.4 | 1.19 | 70.2–77.7 | 3.1 | 2.05 |
| | 2,000 | 73.5 | 1.3 | 1.0 | 1.31 | 70.1–76.0 | 2.4 | 2.23 |
| | 5,000 | 73.3 | 0.8 | 0.6 | 1.23 | 71.1–74.6 | 1.5 | 2.29 |

SD: standard deviation of power between the 50 runs. SD expected from $R$: binomial standard deviation $\sqrt{(P(1-P)/R)}$, where $P$ is the mean power of the 50 runs and $R$ the number of selection simulations. Ratio: SD divided by SD expected from $R$. Each run used twice as many neutral as selection simulations.

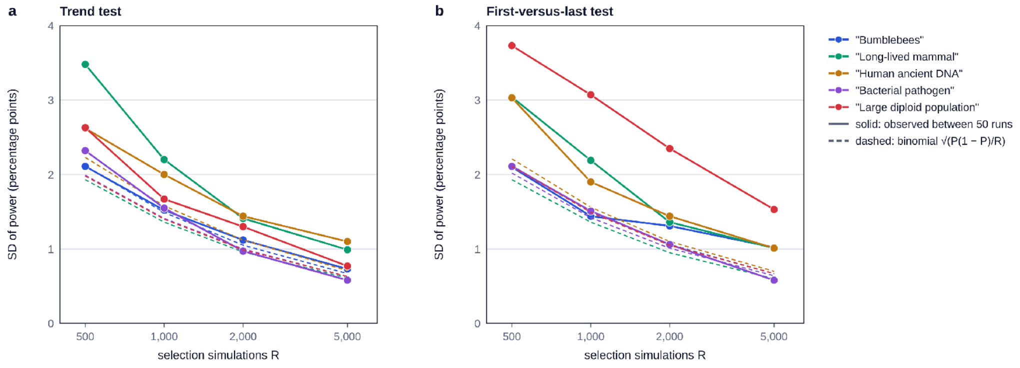


**Figure S3.** Standard deviation of power between 50 runs against the number of selection simulations $R$, for the five example scenarios of Table S7a: (a) trend test; (b) first-versus-last test. Solid lines, observed standard deviation; dashed lines, binomial standard deviation $\sqrt{(P(1 - P)/R)}$, expected if the threshold were fixed (Section S5.1). Each run used twice as many neutral as selection simulations.

### S6 | Rescaling of long time series

#### *S6.1 | Rescaling*

When the first and last sampling dates are more than 2,000 generations apart, SPOT simulates each generation as $\lambda$ real generations (Section 2.6 of the main text). The generation time is multiplied by $\lambda$, so that the generation numbers of the sampling dates are divided by $\lambda$ (and rounded to whole numbers). The breeding numbers $N_f$ and $N_m$, *and* $N_e$ *in each interval, are* divided by $\lambda$. Because the variance of drift per generation is inversely proportional to $N_e$, dividing both the number of generations and $N_e$ by $\lambda$ keeps the total drift over the study period. The selection coefficient becomes $(1 + s)^\lambda - 1$, the advantage accumulated over $\lambda$ generations, and the selection coefficient in males $(1 + s_m)^\lambda - 1$; for small $s$, $(1 + s)^\lambda - 1 \approx \lambda s$, so the product $N_e s$, which sets the strength of selection relative to drift, is kept. The migration rate becomes $1 - (1 - m)^\lambda$, the fraction of breeding adults replaced over $\lambda$ generations. The dominance coefficient $h$, $p_0$ and all sampling and genotyping settings are unchanged. Keeping $h$ unchanged gives heterozygotes the advantage $h[(1 + s)^\lambda - 1]$, close to the advantage they accumulate over λ generations, $(1 + hs)^\lambda - 1$, when $s$ is small.

*Choice of* $\lambda$. SPOT chooses the smallest $\lambda$ for which no more than 2,000 generations are simulated. For example, a population sampled over 17,000 generations is simulated with $\lambda$ = 9, as 1,889 generations. Rescaling is accurate only while the rescaled selection coefficient stays small and the rescaled population stays large (Uricchio & Hernandez, 2014). $\lambda$ is therefore lowered, and more generations are simulated, when needed to keep three conditions: a rescaled selection coefficient of at most 0.025; a smallest rescaled breeding number (over sexes and generations) of at least 200; and a rescaled migration rate of at most 0.5. The first two limits come from the test of Section S6.3. If these conditions would require simulating more than 10,000 generations, $\lambda$ is raised so that 10,000 generations are simulated, and the application marks the result as approximate.

#### *S6.2 | Validation against simulations of every generation*

*Test cases.* We compared rescaled simulations with simulations of every generation. The sampling design had four dates equally spaced over 5,000, 20,000 or 60,000 generations, with 20, 20, 20 and 40 individuals, mean depth 10× at the earlier dates and 15× at the last, per-read error rate 0.005 and minimum depth 3. We combined the three inheritance systems, breeding numbers $N_f = N_m$ = 1,000, 10,000 or 100,000 ($N$ in haploids), $s$ = 0.0001, 0.0005 or 0.002 per generation and $p_0$ = 0.01 or 0.1. Over 60,000 generations, only $N \geq$ 10,000, $s \leq$ 0.0005 and $p_0$ = 0.1 were used, to keep the simulations of every generation affordable, and combinations that SPOT would not rescale ($\lambda$ = 1) were left out. This gave 120 test cases, with $\lambda$ from 3 to 30.

*Procedure.* Each test case was run once with every generation simulated and once with rescaling, each with 1,000 selection and 2,000 neutral simulations and different random seeds, so that the two powers differ by chance even if rescaling is exact. We compared the power of the trend test with the drift-aware threshold, and the run times.

*Results.* In 13 test cases, power was 100% with both. In the other 107, power with rescaling was higher in 50, lower in 53 and equal in four, and the differences had a standard deviation of 2.4 percentage points (1.9 without the haplodiploid test case described below; Table S8; Figure S4). If rescaling had no effect, about 5% of the 107 differences, that is about 5, would exceed two standard errors by chance; 7 did. Without rescaling, the median run time of a calculation grew with the length of the series, from 1.4 s over 5,000 generations to 4.0 s over 20,000 and 15.3 s over 60,000; with rescaling, it stayed between 0.5 and 0.7 s (Figure S4b; run times measured with no other calculation running on the computer). Rescaling thus made calculations up to 28 times faster.

*The largest difference.* One haplodiploid test case (60,000 generations, $N_f = N_m$ = 10,000, $s$ = 0.0005, $p_0$ = 0.1) differed by 14.7 percentage points. We ran it five more times with each method. Mean power was 57% with simulations of every generation and 55% with rescaling, but single runs ranged from 51% to 65% and from 45% to 80%. The difference was therefore not reproduced: over very long series, the threshold can fall where neutral values of $Z$ are sparse, and single runs then vary by more than ten points. Designs whose powers differ by only a few points should therefore be compared with more selection simulations (Section 3.3 of the main text).

*False-positive rate with rescaling.* For nine test cases (the three inheritance systems over 5,000, 20,000 and 60,000 generations, with $\lambda$ = 3, 10 and 30), we set 20 thresholds from rescaled neutral simulations and applied them to 50,000 neutral simulations of every generation. Their mean false-positive rate was 4.7–5.2%, against 10.1–17.6% for the sampling-only threshold.

**Table S8.** Power of the trend test with rescaling minus power with simulations of every generation, by length of the series and inheritance system.

| **Generations** | **Inheritance** | **Test cases** | $\lambda$ | **Mean difference (points)** | **SD of differences (points)** | **Range (points)** | **Speed-up, median (range)** |
|---|---|---|---|---|---|---|---|
| 5,000 | Haploid | 18 | 3 | −0.5 | 0.9 | −3.1 to +1.3 | 2.6 (2.3–2.8) |
| 5,000 | Diploid | 18 | 3 | −0.1 | 1.4 | −2.4 to +2.3 | 2.8 (2.5–2.9) |
| 5,000 | Haplodiploid | 18 | 3 | +0.1 | 1.3 | −3.0 to +2.5 | 2.8 (2.6–2.9) |
| 20,000 | Haploid | 18 | 5, 10 | −0.9 | 2.5 | −7.3 to +2.7 | 7.8 (4.2–8.7) |
| 20,000 | Diploid | 18 | 5, 10 | +0.3 | 1.3 | −2.0 to +3.3 | 8.5 (4.3–9.2) |
| 20,000 | Haplodiploid | 18 | 5, 10 | +0.4 | 1.9 | −2.2 to +6.1 | 7.8 (4.5–9.3) |
| 60,000 | Haploid | 4 | 30 | +1.5 | 1.6 | 0.0 to +3.8 | 25.9 (23.3–26.5) |
| 60,000 | Diploid | 4 | 30 | −1.3 | 1.8 | −3.0 to +0.5 | 26.4 (24.9–27.2) |
| 60,000 | Haplodiploid | 4 | 30 | +4.6 | 7.6 | −2.4 to +14.7 | 25.1 (23.8–28.1) |

Speed-up: run time with every generation simulated divided by run time with rescaling, each measured once with no other calculation running on the computer. Mean difference, standard deviation and range: power of the trend test with rescaling minus power with every generation simulated, in percentage points.

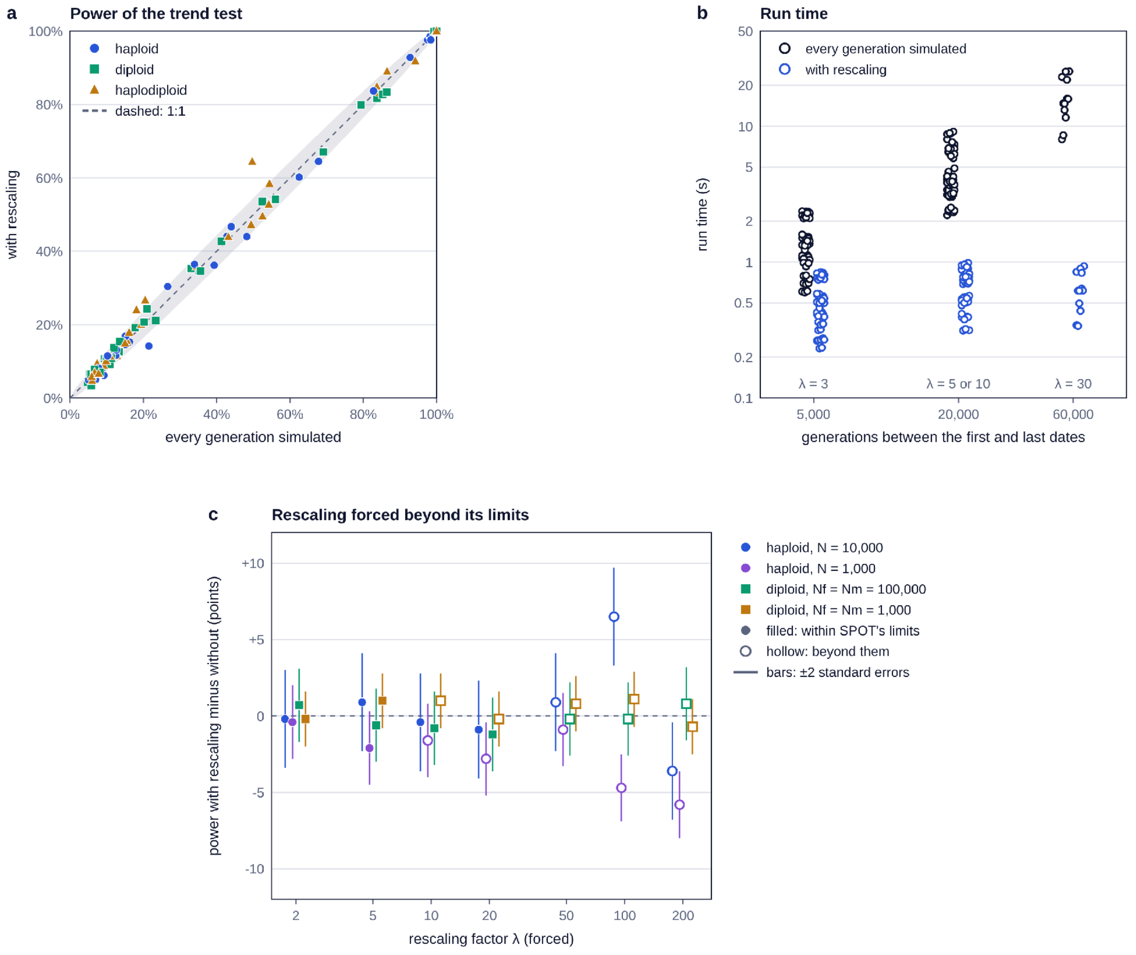


**Figure S4.** Rescaling (Section S6). (a) Power of the trend test with rescaling against power with every generation simulated, for the 120 test cases of Section S6.2, with λ as chosen by SPOT, that is, within the limits of Section S6.1 (colours and shapes, inheritance systems); dashed line, 1:1; grey band, ±2 standard errors of the difference between two powers estimated from 1,000 selection simulations each. (b) Run time of one calculation against the number of generations between the first and last sampling dates, with every generation simulated (black) and with rescaling (blue), for the same test cases; the rescaling factor λ is given under each group; run times measured with no other calculation running on the computer. (c) Rescaling forced beyond SPOT's limits (Section S6.3): power with rescaling minus power with every generation simulated, against the forced rescaling factor λ, in four designs (2,000 selection simulations per power); filled symbols, λ within SPOT's limits; hollow symbols, beyond them; bars, ±2 standard errors; dashed line, no difference.

#### *S6.3 | Limits of rescaling*

To set the limits used in the choice of $\lambda$ (Section S6.1), we forced $\lambda$ from 2 to 200 in four designs and compared the power of the trend test with the power obtained with every generation simulated: haploids with $N = 10{,}000$, $s = 0.0005$ and $p_0 = 0.1$; haploids with $N =$

1,000, $s$ = 0.002 and $p_0$ = 0.1; diploids with $N_f = N_m$ = 100,000, $s$ = 0.0005 and $p_0$ = 0.01; and diploids with $N_f = N_m$ = 1,000, $s$ = 0.0005 and $p_0$ = 0.1. All four had four sampling dates equally spaced over 20,000 generations, with 20, 20, 20 and 40 individuals and the sequencing settings of Section S6.2, and each power was estimated from 2,000 selection simulations. While the rescaled selection coefficient was at most 0.025 and the rescaled breeding numbers at least 200, power differed by at most 2.1 percentage points, within the variation expected between runs (standard error of a difference 0.9–1.6 points). Beyond these values, the largest differences, in the haploid designs, were 4.7–6.5 points, that is, 4–5 standard errors: for example, +6.5 points with a rescaled selection coefficient of 0.051 and 100 rescaled breeding individuals, and −5.8 points with 5 rescaled breeding individuals. In the two diploid designs, power changed by at most 1.2 points at any λ, up to 200 (Figure S4c). The limits are therefore conservative for diploids and protect mainly against the errors seen in haploids; they rest on these four designs only.

#### *S6.4 | Example: a bacterial pathogen*

To illustrate a long series simulated with rescaling, we used the "Bacterial pathogen" example scenario of the application: a haploid bacterium with 100 generations per year ("Years per generation" = 0.01) and $N$ = 100,000 breeding individuals, sampled in 1850, 1900, 1950, 2000 and 2020 (10, 10, 15, 20 and 40 individuals), that is, over 17,000 generations, simulated with $\lambda = 9$ as 1,889 generations. The favoured allele started at $p_0$ = 0.05, with selection from the first date; mean depth was 10× at the earlier dates and 30× at the last, with a per-read error rate of 0.005 and a minimum depth of 3. We computed power for $s$ = 0.00002, 0.00005, 0.0001, 0.0002 and 0.0005 *per generation, with 5,000 selection simulations.*

Power of the trend test *was 8%, 14%, 32%, 73% and over 99% for* these values of $s$ (Figure S5), so 80% power required $s$ between 0.0002 and 0.0005 per generation, about 0.0003 by interpolation, that is, about 0.03% per generation. Even with $N$ = 100,000, the sampling-only threshold was *exceeded by about 7.5% of the neutral* simulations over this many generations. Because power depends on the number of generations rather than on the number of years, a weak selection coefficient can be detected over such a long series.

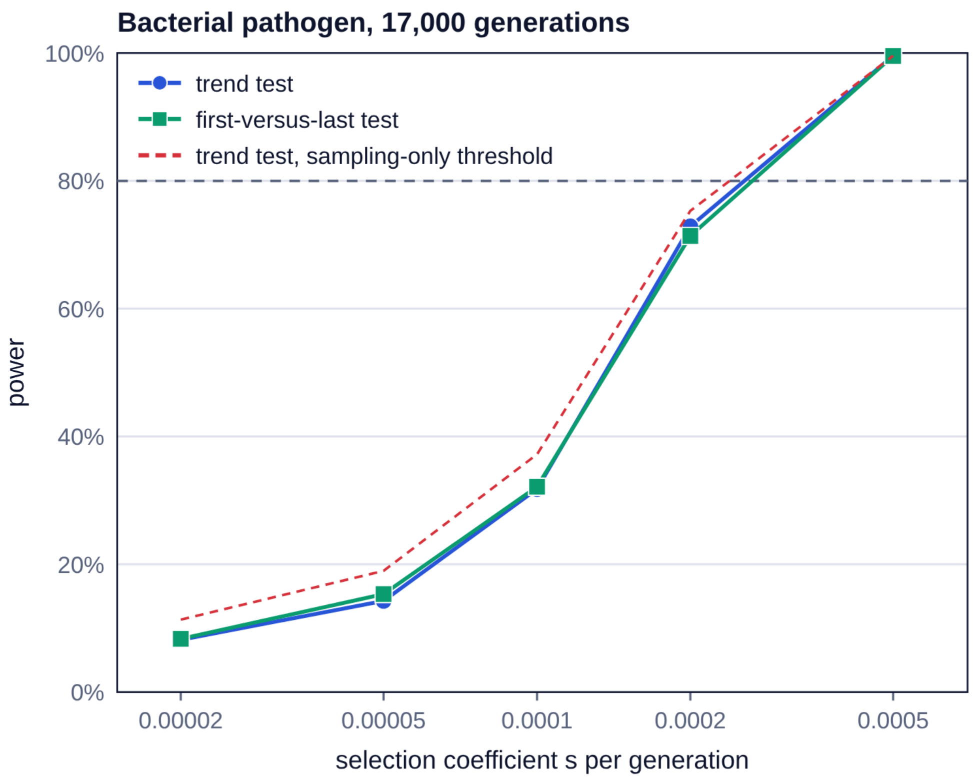


**Figure S5.** Bacterial pathogen example (Section S6.4): power against the selection coefficient per generation, over 17,000 generations simulated with rescaling ($\lambda = 9$), for the trend test and the first-versus-last test with the drift-aware threshold, and for the trend test with the sampling-only threshold (dashed red line). *5,000 selection and 10,000 neutral simulations per value* of *s*; dashed grey line, 80% power.

### S7 | Case studies: tables and figures

This section gives the tables and figures of the two case studies of Section 2.9 of the main text. Their settings are given in Section 2.9 and Table 3 of the main text, and in the legends below.

#### *S7.1 | Museum insect*

**Table S9.** Museum insect case study: power of the trend test (%) for the reference sampling and three ways of adding or reallocating effort (Figure 3d of the main text).

| $N_e$ | $p_0$ | $s$ | Reference (20/20/20/40) | +60 historical (40/40/40/40) | +60 modern (20/20/20/100) | First and last only (50/50) |
|---|---|---|---|---|---|---|
| 1,500 | 0.01 | 0.05 | 67.0 | 68.8 (+1.7) | 69.1 (+2.1) | 66.6 (−0.5) |
| 1,500 | 0.1 | 0.02 | 55.3 | 59.9 (+4.6) | 60.9 (+5.6) | 60.6 (+5.3) |
| 1,500 | 0.3 | 0.015 | 55.1 | 57.3 (+2.2) | 59.9 (+4.8) | 61.6 (+6.5) |
| 100,000 | 0.01 | 0.035 | 63.0 | 74.0 (+11.1) | 84.4 (+21.5) | 78.6 (+15.7) |
| 100,000 | 0.1 | 0.015 | 55.9 | 67.4 (+11.5) | 72.0 (+16.1) | 68.3 (+12.4) |

| $N_e$ | $p_0$ | $s$ | Reference (20/20/20/40) | +60 historical (40/40/40/40) | +60 modern (20/20/20/100) | First and last only (50/50) |
|---|---|---|---|---|---|---|
| 100,000 | 0.3 | 0.012 | 63.3 | 74.5 (+11.2) | 76.6 (+13.3) | 77.1 (+13.8) |

Reference: 20, 20, 20 and 40 females sampled in 1935, 1955, 1975 and 2025. +60 historical: 40 females at each historical date. +60 modern: 100 females in 2025. First and last only: the same 100 females, 50 in 1935 and 50 in 2025. In brackets: change from the reference sampling, in percentage points. Haplodiploid population with $N_e$ = 1,500 ($N_f = N_m$ = 1,000) or 100,000 ($N_f = N_m$ = 66,667); for each $N_e$ and $p_0$, $s$ was chosen so that the reference sampling had intermediate power (55–67%). Other settings as in the reference case (Table 3 of the main text): mean depth 10× at the historical dates and 15× in 2025, per-read error rate 0.005, minimum depth 3, $h$ = 0.5, unrelated individuals, selection from the first date; trend test with the drift-aware threshold, $\alpha$ = 0.05, one locus. 5,000 selection and 10,000 neutral simulations per design.

### *S7.2 | Human ancient DNA*

**Table S10.** Human ancient DNA case study: power with the same number of reads (100) per ancient date, spread over more or fewer individuals (Figure S6b).

| Individuals per ancient date | Mean depth | Individuals without a genotype (%) | Heterozygotes called correctly (%) | Power, $s$ = 0.005 (trend / first-versus-last) | Power, $s$ = 0.0075 (trend / first-versus-last) |
|---|---|---|---|---|---|
| 100 | 1× | 30.7 | 46.6 | 47.6 / 43.1 | 79.1 / 74.1 |
| 50 | 2× | 9.7 | 66.5 | 46.8 / 40.8 | 77.8 / 71.2 |
| 25 | 4× | 1.0 | 88.0 | 44.6 / 38.8 | 76.1 / 68.0 |
| 20 | 5× | 0.3 | 92.6 | 42.6 / 35.3 | 73.2 / 64.2 |
| 10 | 10× | 0.0 | 98.7 | 37.1 / 29.6 | 66.4 / 53.1 |

Each design has the same number of reads at the locus per ancient date (number of individuals × mean depth = 100). Individuals without a genotype: proportion of all sampled individuals, ancient and present-day, with fewer reads than the minimum depth (here 1 read), who receive no genotype (Section S1.8). Heterozygotes called correctly: proportion of the true heterozygotes that received a genotype (ancient and present-day pooled) and were called heterozygous; at low depth, many heterozygotes are called homozygous because all their reads show the same allele. Both proportions for $s$ = 0.005 (almost identical for $s$ = 0.0075). Settings of the "Human ancient DNA" example scenario (Table 3 of the main text; Table S7a): diploid, $N_f = N_m$ = 5,000 ($N_e$ = 10,000), 28 years per generation, samples in 7000, 5000, 3000 and 1000 BCE and 1000 and 2000 CE (321 generations), $p_0$ = 0.1, $h$ = 0.5, selection from the first date, per-read error rate 0.01, minimum depth 1; the present-day sample is 100 individuals at 20× in all designs. Trend test with the drift-aware threshold, $\alpha$ = 0.05, one locus; 5,000 selection and 10,000 neutral simulations per design.

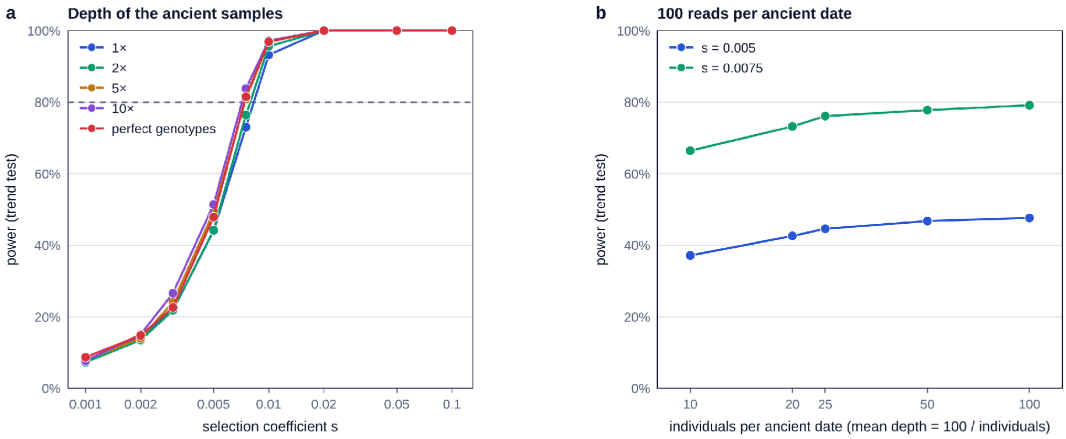


**Figure S6.** Human ancient DNA case study. (a) Power of the trend test against the selection coefficient, with the ancient samples sequenced at 1×, 2×, 5× or 10×, or with perfect genotypes; s from 0.001 to 0.1; 1,000 selection and 2,000 neutral simulations per design. (b) Power with the same 100 reads per ancient date spread over 100 individuals at 1× to 10 individuals at 10× (s = 0.005 and 0.0075); 5,000 selection and 10,000 neutral simulations per design. Other settings as in Table S10.

### S7.3 | Pseudo-haploid calls

In human ancient DNA, genotypes are often replaced by pseudo-haploid calls: for each individual with at least one read at the locus, one read is drawn at random and counts as one gene copy; an individual without a read gives nothing. We simulated the human ancient DNA case study with the ancient samples at 1×, 2×, 5× or 10× and the present-day sample at 20×, and analysed the same reads in two ways: with SPOT's genotype calls and with pseudo-haploid calls for all individuals. For each analysis, the threshold was set from 10,000 neutral simulations analysed the same way, and power was measured on 5,000 selection simulations. With pseudo-haploid calls, power was 6–12 points lower, and it increased less with depth: with $s$ = 0.0075, from 66% at 1× to 72% at 10×, against 73% to 84% with SPOT's genotype calls. This is because a pseudo-haploid call uses one read per individual, however many reads the individual has. At 1×, about a third of the individuals have no read at the locus and give nothing; at 5×, almost all have at least one read. Deeper sequencing therefore helps only until nearly every individual has a read, after which the additional reads are not used.

### S8 | Reproducibility

*Random seed.* Each calculation of SPOT starts from a random seed ("Random seed" in the application). With the same seed and the same settings, SPOT gives exactly the same results; with another seed, the results differ by the variation between runs described in Section S5.

*Code.* The application (index.html) and its simulation code, which is also distributed as a separate file (sim.js) that runs in Node.js, are in the SPOT repository on GitHub and in its archived version (Data Availability Statement). All analyses of this article were run with sim.js, with the scripts listed below, which are in the folder paper of the repository. Each script sets its own random seeds, so that it reproduces the results exactly, and writes a tab-separated results file; the figures are drawn from these files by Python scripts that use only the standard library.

*False-positive rate (Section S2).* false_positive_rate.js, run with the letters A, B and C, gives Tables S1, S2 and S3 (files fpr_A.tsv, fpr_B.tsv and fpr_C.tsv); with the letter G, it measures the threshold from a normal distribution fitted to 2,000 neutral simulations (fpr_C_gaussian.tsv). normal_fit_rare.js measures the threshold used beyond 1,000 loci (Section S1.12); nodrift_example.js computes the *example without drift of Section S1.11.*

*Agreement with theory (Section S3).* theory_spot.js runs SPOT (theory_spot.json), and theory_check.py calculates the theoretical values independently of SPOT's code and compares them (theory_power.tsv and theory_drift.tsv); sibling_path_check.js compares the two sampling procedures of Section S3.4 (sibling_path_check.tsv). migration_diversity_check.js compares the loss of diversity with migration to Equation (7) (migration_diversity.tsv).

*Comparison with SLiM (Section S4).* The SLiM model is in slim_check.slim, run by slim_runs.sh and slim_runs_v2.sh. The first run of each scenario used seed 12345, the four further runs of 2,000 neutral simulations used for the false-positive rates seeds 2, 3, 4 and 5, and runs 2–20 seeds 1,000 + $k$ (scenarios with selection) and 2,000 + $k$ (scenarios without selection) for run $k$. slim_compare.js compares the allele frequencies (SPOT seeds 100 + $j$ and 200 + $j$ for scenario $j$); slim_compare_v2.js compares power (SPOT seeds 10,000 + 100$j$ + $k$ and 20,000 + 100$j$ + $k$ for the selection and neutral simulations of run $k$ of scenario $j$) and false-positive rates (seeds 30,000 + 100$q$ + $i$, $i$ = 0–9, for the 20,000 simulations that set the threshold, and 40,000 + 100$q$ + $i$, $i$ = 0–4, for the 10,000 independent ones, for scenario without selection *q*); slim_power_spot_threshold.js computes power with SPOT's threshold (threshold from seeds 50,000 + 100*j* + *i*, *i* = 0–9); slim_regression.py fits the regressions. Scenarios are numbered from 0 in the order of Tables S5 and S6.

*Precision (Section S5).* monte_carlo_precision.js runs the five example scenarios 50 times, with seed 5,000 + 97*i* for run i = 0–49 (mc_precision.tsv); speed.js measures the run times (speed.tsv).

*Rescaling (Section S6).* rescale_validation.js compares rescaled simulations with simulations of every generation (rescale_results.tsv) and, run with the argument times, measures the run times with no other calculation running (rescale_times.tsv); rescale_hd_repeat.js repeats the haplodiploid test case that differed by 14.7 points; rescale_fpr.js measures the false-positive rate of thresholds set from rescaled neutral simulations; rescale_limits.js forces the rescaling factor (rescale_limits.tsv); worked_examples.js, run with the letter E, computes the bacterial pathogen example.

*Case studies (Sections 2.9 and 3.4 of the main text; Sections S6.4 and S7).* worked_examples.js computes the case studies, with one letter per analysis: A, power against *s* in the museum insect case study (Figure 3a; bee_s_p0.tsv); B, where to add effort (Figure 3d; Table S9; bee_allocation.tsv); C, power against $N_e$ (Figure 3b,c; bee_ne.tsv); E, the bacterial pathogen (Section S6.4; Figure S5; pathogen.tsv); F and G, human ancient DNA (Figure S6; Table S10; human_depth.tsv and human_budget.tsv). depth_case.js computes the effect of sequencing depth (Figure 3e; depth_case.tsv). headline_runs.js repeats the reference case 20 times (Section 3.4 of the main text; headline_runs.tsv). pseudohaploid_check.js compares SPOT's genotype calls with pseudo-haploid calls in the human ancient DNA case study (Section S7.3; pseudohaploid.tsv).

### References


Armitage, P. (1955). Tests for linear trends in proportions and frequencies. *Biometrics*, *11(3)*, 375–386. https://doi.org/10.2307/3001775

Cochran, W. G. (1954). Some methods for strengthening the common $\chi^2$ tests. *Biometrics*, *10(4)*, 417–451. https://doi.org/10.2307/3001616

Crow, J. F., & Kimura, M. (1970). *An introduction to population genetics theory*. Harper & Row.

Haller, B. C., Ralph, P. L., & Messer, P. W. (2026). SLiM 5: Eco-evolutionary simulations across multiple chromosomes and full genomes. *Molecular Biology and Evolution*, *43(1)*, msaf313. https://doi.org/10.1093/molbev/msaf313

Hedrick, P. W., & Parker, J. D. (1997). Evolutionary genetics and genetic variation of haplodiploids and X-linked genes. *Annual Review of Ecology and Systematics*, *28(1)*, 55–83. https://doi.org/10.1146/annurev.ecolsys.28.1.55

Nielsen, R., Paul, J. S., Albrechtsen, A., & Song, Y. S. (2011). Genotype and SNP calling from next-generation sequencing data. *Nature Reviews Genetics*, *12(6)*, 443–451. https://doi.org/10.1038/nrg2986

Uricchio, L. H., & Hernandez, R. D. (2014). Robust forward simulations of recurrent hitchhiking. *Genetics*, *197(1)*, 221–236. https://doi.org/10.1534/genetics.113.156935

Wright, S. (1931). Evolution in Mendelian populations. *Genetics*, *16(2)*, 97–159. https://doi.org/10.1093/genetics/16.2.97
